# Evolution of two-component quorum sensing systems

**DOI:** 10.1101/2020.11.15.381053

**Authors:** Marina Giannakara, Vassiliki Lila Koumandou

## Abstract

Quorum sensing (QS) is a cell-to-cell communication system that enables bacteria to coordinate their gene expression depending on their population density, via the detection of small molecules called autoinducers. In this way bacteria can act collectively to initiate processes like bioluminescence, virulence and biofilm formation. Autoinducers are detected by receptors, some of which are part of Two Component Signal Transduction Systems (TCS), which comprise of a (usually membrane-bound) sensor histidine kinase (HK) and a cognate response regulator (RR). Different QS systems are used by different bacterial taxa, and their relative evolutionary relationships have not been extensively studied. To address this, we used the KEGG database to identify all the QS HKs and RRs that are part of TCS and examined their conservation across microbial taxa. We compared the combinations of the highly conserved domains in the different families of receptors and response regulators using the SMART and KEGG databases, and we also carried out phylogenetic analyses for each family, and all families together. The distribution of the different QS systems across taxa, indicates flexibility in HK/RR pairing, and highlights the need for further study of the most abundant systems. For both the QS receptors and the response regulators, our analysis indicates close evolutionary relationships between certain families, highlighting a common evolutionary history which can inform future applications, such as the design of novel inhibitors for pathogenic QS systems.

## Introduction

### The mechanism and importance of quorum sensing

Quorum sensing is a mechanism responsible for regulating a variety of group behaviors and interactions in bacteria, including pathogenicity, biofilm formation, mobility, sporulation, conjugal plasmid transfer, bioluminescence, resistance to antibiotics, and antibiotics production (Bandara et al., 2012; Ng and Bassler, 2009). All quorum sensing systems, despite in their molecular mechanisms and regulatory components, regulate their target genes in a population-density dependent manner. An extracellular molecule (autoinducer, AI) is secreted by a group of bacteria and when the number of the bacteria rises, so does the number of these molecules produced in the surrounding area. When the density of the AIs overcomes a specific threshold, they bind to a receptor protein, either on the membrane or in the cytoplasm of the cell. The detection of the AIs activates a signal transduction cascade, altering the gene expression of the cells all over the population (Ng and Bassler, 2009).

The quorum sensing mechanism is different between Gram-positive and Gram-negative bacteria. Gram-positive bacteria use modified oligopeptides as AIs (autoinducing peptides, AIPs), which bind to membrane-bound receptors -Histidine Kinases (HKs) -when they exceed a density threshold. This leads to the phosphorylation of the HK protein and its cognate response regulator (RR), which then regulates the target genes. The AIPs can also be transported into the cell and interact with cytoplasmic AIP receptors, which act as transcription regulators to regulate gene expression (Ng and Bassler, 2009; Rutherford and Bassler, 2012). Gram-negative bacteria use AHLs (acyl-homoserine-lactones) or SAM-products (S-adenosylmethionine)(Papenfort and Bassler, 2016). A second group of AIs are DPD (4,5-dihydroxy-2,3-pentanedione) derived molecules, collectively known as AI-2 (Bandara et al., 2012). Those small molecules can diffuse through the cell membrane and bind to LuxR-type cytoplasmic receptors, which function as transcription factors. However, in some Gram negative bacteria the AIs are detected by Histidine Kinases and the process is similar to that in the Gram positive bacteria (Rutherford and Bassler, 2012). Other inducer molecules taking part in quorum sensing include protein molecules that are not described as AIPs, pheromones, sulfate and phosphate, fucose, oxygen, quinolones, γ-aminobutiric acid (https://www.genome.jp/kegg-bin/show_pathway?map02024) or even adrenaline and noradrenaline produced by the gut bacteria in humans, known as AI-3 (Reading and Sperandio, 2006).

Since quorum sensing affects many bacterial processes, it is an interesting subject of study which can have numerous applications, most important of which is the control of pathogenicity via QS inhibition, given the increasing spread of antibiotic-resistant bacteria. It has been suggested that targeting QS imposes less selective pressure, since it does not kill bacteria but only hinders the production of virulence factors (Whiteley et al., 2017). QS inhibition (Quorum Quenching, QQ) can be applied to fields such as hospital infections, phytopathogen control in agriculture and for producing new preservatives in the food industry (Sreenivasulu, 2015; Tiwari et al., 2016). The gene products involved in QS mechanisms are consequently possible targets in new antimicrobial strategies and the investigation of their evolutionary relationships can contribute to that direction (Lerat and Moran, 2004).

### Two-Component System types

The Histidine Kinases that contribute to QS in both Gram positive and negative bacteria are part of the Two Component Signal Transduction (or Component) Systems (TCSs). TCS is the main mechanism in bacteria for responding to environmental stimuli and prevails across the entire bacterial kingdom (Capra and Laub, 2012). Although there is a wide variety in TCS mechanisms, they all fall into two main categories: The canonical and the multi-step TCS.

### Canonical TCS: His-Asp

A typical TCS comprises two proteins: a transmembrane Histidine Kinase (HK) and a Response Regulator (RR). The HK is a dimeric protein which includes 3 domains: The Sensor, the HisKA (Histidine Kinase domain or Dimerization domain) and the HATPase (ATP kinase binding domain). The sensor domain binds to the AI and this interaction results in an ATP-dependent autophosphorylation of the conserved histidine of HisKA. The phosphoryl group is then transferred to a protein called Response Regulator (RR). The RR contains the Response Regulator domain (or REC or Receiver domain) in the N-terminal region and the Effector domain in the C-terminus (Jacob et al., 2014; Kaserer and West, 2010). The first domain includes the conserved aspartate residue, which is phosphorylated (Kaserer and West, 2010). This leads to conformational changes in the structure of the RR (West and Stock, 2001), which is responsible for controlling quorum sensing-associated behaviors through the Effector domain either via protein-protein interactions or by binding to DNA for gene regulation (Mascher et al., 2006).

### Multi-step TCS: His-Asp-Asp

A more complicated TCS mechanism usually includes a hybrid HK (HHK), a histidine phosphotransferase (HPt) and its cognate RR. The structure of the HHK is similar to HK but additionally includes a conserved Asp residue in the C-terminus, namely a REC domain. The binding of the signaling molecule induces the autophosorylation of the His of the HisKA domain and then the phosphoryl-group is transferred intramolecularly to the Asp residue of the HHK. Then the phosphotransfer continues to the HPt protein and from there to the RR (Capra and Laub, 2012; Kaserer and West, 2010).

### Categories of HKs and RRs

The classification of the HKs and the RRs into groups varies and depends on the comparison criteria. In the KEGG database the HKs and their cognate RRs are divided into separate families (Supplementary Fig. S1); however, the criterion for this classification is not obvious. In previous studies, the HKs and RRs were classified based on their conserved domains. For example, the HKs were divided into 11 subfamilies, based on multiple alignment of 348 HKs and comparison of the “Homology boxes”, i.e. groups of highly conserved amino acids, which are assumed to be important in substrate binding, catalysis or the structure of the protein: the H- and X-boxes are parts of the HisKA domain, and the N-,D-F-and G-boxes are part of the HATPase_c domain; the D-box is also part of the REC domain (Grebe and Stock, 1999). Alterntively, given the structure of the proteins and more specifically their domain organization, the HKs can be divided into 4 groups: 3 groups with proteins containing the intracellular domains PAS, HAMP or GAF, respectively, and a group with none of these domains (Capra and Laub, 2012). In a more recent study, the proteins were separated into 5 types (Types I-IV and the chemosensor HKs group, CheA) based on the inferred phylogenetic tree using the whole genome of 22 bacterial species and 4 archaea. More specifically, the criterion for their division was the H-box (HisKA) and its secondary structure, and the kinase domain (ATPase) (Forst and Kim, 2015).

Based on the DNA-binding domains (Effector domain) of RRs, they were classified into three subfamilies: OmpR, Fix J and NtrC (Hakenbeck and Stock, 1996), or more recently into 5 groups (Capra and Laub, 2012). By comparing the Receiver domains (REC), the RRs were classified into 8 subfamilies (R_A_-R_H_) (Grebe and Stock, 1999), however, it has been proposed that the comparison of the RRs for their classification should be based only on the Effector domain, due to the fact that the Receiver domains of different RRs are highly conserved and therefore not very informative for classification (Galperin, 2006).

### Evolution of TCS

It is suggested that the more variable the environment, the higher the number of the TCS of the organisms living in it. The genes of the HKs and their cognate RRs are usually found on the same operon. The various TCS pathways are the result of lateral gene transfer and/or duplication. The duplication can happen to all of the genes of the operon or to one of them; in the second case, the resulting TCS pathway consists of more than one HK and responds to more than one extracellular signal, or it includes more than one RR and gives multiple responses for a specific signal. Also, due to the modular nature of HKs, domain shuffling can lead to new HKs and the fusion of the HK and RR genes of an operon can create hybrid HKs. The function of TCS pathways relies on molecular recognition and consequently the amino acids have extensively co-evolved as a means to prevent the disruption of signaling by mutations. Co-evolution also prevents crosstalk with other pathways. Concerning the origin of HKs, it has been suggested that they come from ATPases of the GHKL superfamily: Hsp90, the mismatch repair protein MutL or type II topoisomerases (Capra and Laub, 2012; Koretke et al., 2000). On the other hand, the origin of the RRs remains unclear beyond a general structural similarity to P-loop NTPases (Koretke et al., 2000). The evolution of RRs can be the result of changes in the DNA-binding or RNA-polymerase interaction sites; it includes lateral gene transfer and gene duplication, while domain shuffling and rearrangement have also led to new forms of RRs (Capra and Laub, 2012).

### Evolution of QS

Most of the phylogenetic studies of QS have mainly focused on the produced AIs or on specific pathways, mainly LuxI/LuxR and LuxS/LuxQ. A phylogenetic analysis for the proteins LuxI, LuxR and LuxS, showed that these proteins are ancient in many bacterial species and that they appeared very early in the evolutionary path of the bacteria. It was also found that in most cases the gene pairs of the inducer and its cognate response regulator are located next to each other on the chromosomes and they preserve their pairwise function, a sign of common evolutionary history (Lerat and Moran, 2004). The existence of homologous proteins in some genomes is representative of horizontal transfer and duplication of the genes, both of which can alter the regulation of different gene targets (Lerat and Moran, 2004). In a study of the ComQXPA pathway, the phylogenetic tree of HK proteins in 60 firmicute genomes containing the comQXPA locus clustered all the ComP proteins from various organisms together, instead of gathering them with HKs of the same organism. Therefore, it concluded that ComP evolved from the other HKs before the appearance of the modern bacterial species and thus the ComQXPA pathways have an ancient origin (Dogsa et al., 2014).

In a study of the Vibrionaceae family, it was found that the various inducers (AHLs) do not show any correlation with the species’ geographical distribution. This result indicates that AHLs had a worldwide distribution throughout their evolution, since there was no identification of specific AHL to the environment of the organism which produces it (Rasmussen et al., 2014). Another survey, focusing on thermophilic bacteria, showed that they use AIs-2 for QS communication. The phylogenetic tree comparison of LuxS and the 16S rRNA of thermophiles and mesophiles show that LuxS (which produces the autoinducer) of mesophilic bacteria may originate from thermophiles. Also, the LuxS proteins of thermophilic bacteria within a phylum were evolutionarily closer compared to LuxS of different phyla (Kaur et al., 2018).

In this study we examine the evolutionary relationships between different TCS QS systems. To achieve this, we conducted phylogenetic and comparative analysis of the components of these pathways. We used the KEGG database to identify all of the QS proteins that are also part of the TCS, and we collected their amino acid sequences from various bacterial species. The KEGG and SMART databases were used to compare the combinations of highly conserved domains in HKs and RRs. Phylogenetic analysis was based on Maximum Likelihood methods.

## Methods

### Distribution of QS proteins

We used the **KEGG Database** (Kyoto Encyclopedia of Genes and Genomes, https://www.kegg.jp/ or https://www.genome.jp/kegg/), to select the amino acid sequences used in this study. First, the Two Component System protein types that are also part of the Quorum Sensing system were found using the pathway maps of the KEGG PATHWAY database (maps ko02020 and ko02024 respectively). Each protein type is attributed to a specific orthology group accession number. By entering each identifier in the KEGG ORTHOLOGY database section, we were able to gather information about the amino acid sequences of each protein type in different bacterial organisms (“Genes” section) and the lineages in which each protein type is present (“Taxonomy”). We checked for the presence of the protein subunits of different QS pathways in selected groups of bacteria, including various Phyla, Classes and Genera, as shown in Supplementary Table S1.

### Conserved domains

One amino acid sequence for each protein type was chosen randomly to identify the conserved domains on it, by searching in the “motifs” section in the KEGG GENES database. We also used SMART (Simple Modular Architecture Research Tool, http://smart.embl-heidelberg.de/) to identify additional domains in our selected amino acid sequences not shown in KEGG. Both SMART and KEGG also give the E-value of each conserved domain; the domains that were finally taken into consideration for our research, were those with an E-value of at least 10^−5^ which were also mentioned as “Confidently predicted domains” in SMART. In general, most of the domains were the same in the two databases. Next, using the Pfam database (https://pfam.xfam.org/), we found the groups (clans) in which these domains are classified (Supplementary Table S2) as well as information about their function. For further information we used the Conserved Domain Database (CDD) in NCBI (https://www.ncbi.nlm.nih.gov/Structure/cdd/cdd.shtml).

### Taxa and families representation

For the phylogenetic analysis, the number of the collected amino acid sequences for each protein type was kept as low as possible so as to have a manageable amount of data but at the same time include a representative portion of the different taxa and families which include each protein type. As a result, the average number of the amino acid sequences for each protein type was about 20. Supplementary Tables S3A and S3B show the representation percentages of the taxa, families and species out of the total available ones in the KEGG database. Accession numbers and abbreviations for all sequences used are in the Supplementary dataset.

### Phylogenetic analysis

Multiple alignments of the amino acid sequences were created using the MAFFT (Multiple Alignment using Fast Fourier Transform, https://www.ebi.ac.uk/Tools/msa/mafft/) (Madeira et al., 2019) and MUSCLE (MUltiple Sequence Comparison by Log-Expectation, https://www.ebi.ac.uk/Tools/msa/muscle/) (Edgar, 2004), alignment programs, with default settings, for:

- each HK family
- all the HKs
- each RR family
- all the RRs

The substitution model for each multiple alignment was inferred via the ProtTest3.4 program (https://github.com/ddarriba/prottest3/releases) (Pearson, 2013). In order to compute the likelihood scores, the following settings were used:

- Substitution matrices to test: VT, Blosum62, JTT, Dayhoff, WAG
- Rate variation: I, G, I+G
- Amino acid frequencies: unchecked empirical

The results were evaluated by the Akaike information criterion (AIC). More specifically, the substitution model for the MUSCLE multiple alignments of the HKs and RRs was VT and for the MAFFT alignments was VT for HKs and WAG for RRs.

Phylogenetic reconstruction for each protein family was done with PhyML at http://www.phylogeny.fr/ (also available at https://ngphylogeny.fr/) (Dereeper et al., 2008). We used the “A la Carte” function, with the following settings: “remove gaps in alignment” (alignment curation), Maximum Likelihood method (PhyML), SH-like Approximate Likelihood-Ratio Test (aLRT) as the statistical test for branch support and 100 bootstraps. For each multiple alignment we chose the WAG substitution model as the best one according to the ProtTest results (WAG, JTT and Dayhoff were the given options in the PhyML settings).

The phylogenetic trees for all the protein families together (2 trees: HKs and RRs) were constructed in the CIPRES Science Gateway (https://www.phylo.org/portal2/login.action), using RAxML (Randomized Axelerated Maximum Likelihood): *RAxML-HPC BlackBox (8*.*2*.*12) -Phylogenetic tree inference using maximum likelihood/rapid bootstrapping on XSEDE* (Stamatakis, 2014). Before entering the multiple alignments into the program, we removed any positions not conserved in >50% of the sequences. For this purpose, we used Jalview (https://www.jalview.org/) to detect those positions and then the Mesquite program to do the trimming (http://www.mesquiteproject.org/). The proper substitution model was chosen for each multiple alignment and the bootstrap number was set to 100.

The multiple alignment for all the HK proteins consisted of 347 sequences and had a total length of 2410 amino acids using MAFFT or 1919 using MUSCLE. In both alignments, there were several clusters of conserved residues mixed with gaps or non-conserved residues towards the N-terminal region of the alignment (41-867) (34-1073, 1295-1781) and near the C-terminal part (949-1411). The trimmed MAFFT alignment had 520 amino acids (Supplementary Fig. S2), while the trimmed MUSCLE alignment had 455 amino acids (Supplementary Fig. S3). The multiple alignment for all the RR proteins consisted of 293 sequences and had a total length of 751 amino acids using MAFFT or 670 using MUSCLE. Both the MUSCLE and MAFFT alignments contained conserved clusters alternating with gaps or non-conserved residues (65-224, 287-528). Trimming resulted in an alignment of 229 amino acids using MAFFT (Supplementary Fig. S4), and 274 using MUSCLE (Supplementary Fig. S5).

All the phylogenetic trees were visualized with the Figtree program (https://github.com/rambaut/figtree/releases). The results were saved as PNG images and were further edited with Microsoft Draw. Nodes were labelled as white, grey or black bullets, if their bootstrap value was 50%-80%, 80%-95% and 95%-100% respectively. The raw and masked alignments, as well as the phylogenetic trees in nexus format are available in the Supplementary dataset. The alignments for each family and all families together are available in the files “HK alignments” and “RR alignments”. The alignments of all HKs and all RRs are presented as they were trimmed in Mesquite: in the added taxon, “C” and “I” represent the removed and included positions. The trees in nexus format are available in the “HKs trees” and “RRs trees” files.

## Results

### Distributions of QS proteins

The distribution of each protein type across bacterial taxa is demonstrated in Fig. 1; bacterial groups are colored based on (Jun et al., 2010). All pathways are fully present in at least one bacterial group, but many groups of bacteria have some of the subunits of each pathway. All Lux pathways are fully present in γ-proteobacteria. The Lux system contains a pathway of HKs and an RR, and the cytoplasmic Lsr proteins; both pathways are activated by the AI produced by LuxS. Deinococci, actinobacteria, clostridia, bacilli, α-and γ-proteobacteria, and Spirochaetes possess most or all proteins of the Lsr protein pathway. The UvrY pathways (BarA/UvrY and SdiA) are fully present in γ-proteobacteria. Like in the Lux pathway, there are two different types of receptors, an HK (BarA) and a cytoplasmic protein (SdiA). Several proteins of one or both of the pathways are also found in α-β-and δ-proteobacteria, chlorobi, bacilli, *Nitrospira*, Chrysiogenetes and Spirochaetes. Concerning the NarL pathway, γ-proteobacteria have all proteins of the pathway; α-and β-proteobacteria, chloroflexi, cyanobacteria, actinobacteria, clostridia and bacilli have one of the receptors. Only γ-proteobacteria possess all proteins of the Rcs pathway; bacteroidetes and β-proteobacteria contain the HK and RR of the pathway. All three proteins of the Fus pathway are in γ-proteobacteria; however, the HK/RR pair is found also in α- and β-proteobacteria, bacteroidetes, deinococci, actinobacteria and Spirochaetes. All proteins of the Com(XQPASK) pathway are found in bacilli and most proteins are found in clostridia. Also, the HK/RR pair ComP/ComA is present in deinococci. The proteins of Com(ABDEX) are all found in bacilli, whereas the HK/RR pair ComD/ComE is present in clostridia. The Agr pathway (both AgrC/AgrA and SaeS/SaeR) is fully present in bacilli, while α-proteobacteria and clostridia contain the Agr proteins only. All the proteins of the Nis pathway are present in both clostridia and bacilli, while bacteroidetes, chloroflexi and actinobacteria contain all but two of the proteins. Other bacterial groups contain parts of the pathway but not the full sensing system (they lack either one or both the HK and RR). Bacteroidetes, actinobacteria, clostridia and bacilli contain both the CiaH and CiaR proteins. Concerning the Qse pathway, γ-proteobacteria contain all of the proteins (QseC/QseB and QseE(GlrK)/QseF(GlrR)). QseC/QseB are present in bacteroidetes, actinobacteria and aquificae. QseE(GlrK)/QseF(GlrR) are found in β- and δ-proteobacteria and *Nitrospira*. No bacterial group contains all the proteins of the Rpf pathway: Nitrospira, α- and β-proteobacteria contain the HK/RR pair RpfC/RpfG. All three proteins of the DesR pathway are present in α-,β- and γ-proteobacteria, actinobacteria and bacilli. For details of all other bacterial groups which contain only certain proteins of each pathway, please refer to Fig. 1.

**Figure 1:**
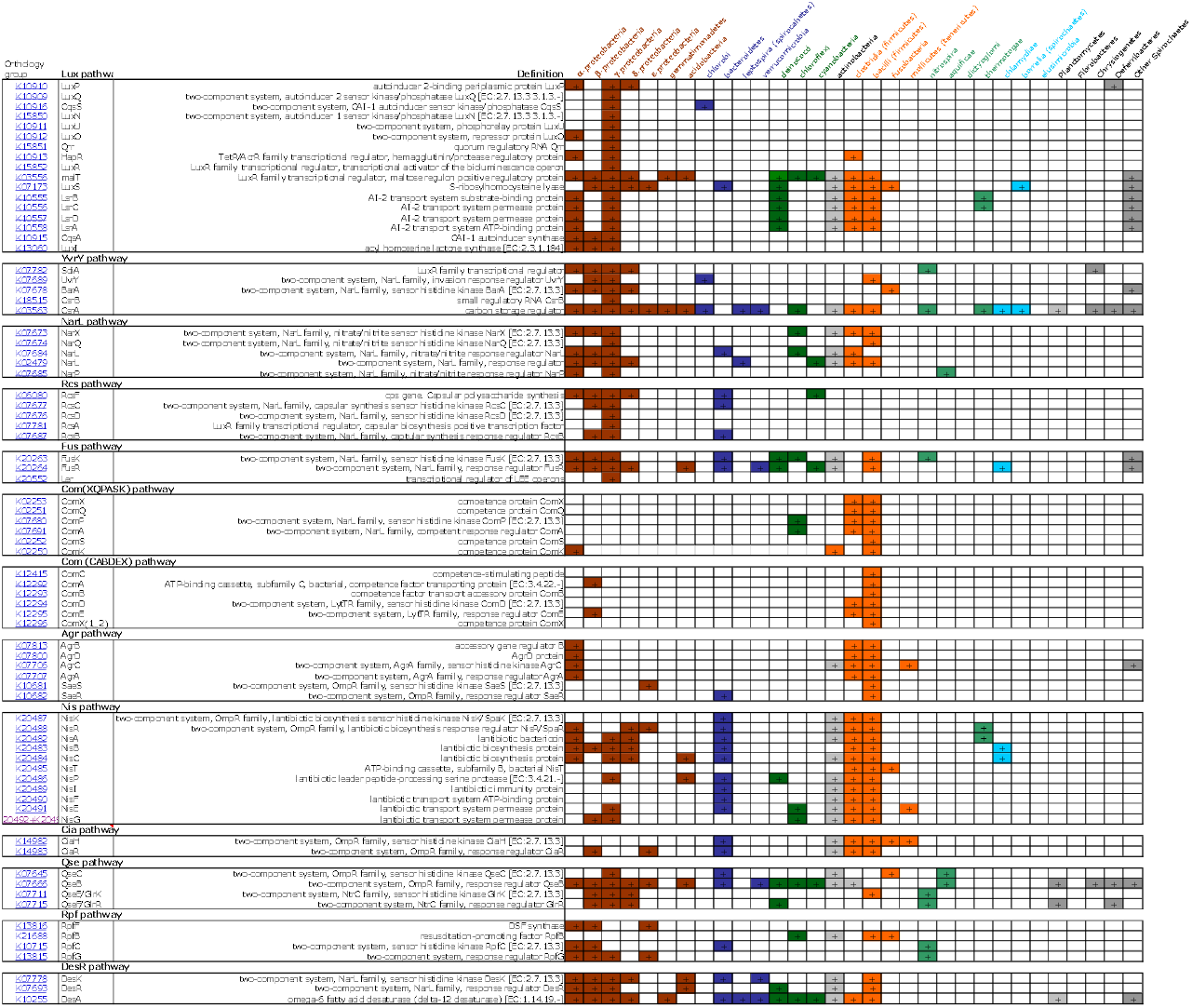
Presence of the protein subunits of different QS pathways in selected groups of bacteria. Colored sectors indicate presence, empty sectors indicate absence, based on the KEGG taxonomy database for each orthology group. “Other” Spirochaetes refer toTreponema, Spirochaeta, Leptospira and Salinispira.

Fig. 1 shows that the majority of the complete pathways are found in α-, β- and γ-proteobacteria, clostridia and bacilli. A significant number of proteins are also found in δ- and ε-proteobacteria, bacteroidetes and actinobacteria. Also, based on the number of bacterial groups which contain proteins of each pathway, the most widespread proteins are those of the UvrY, DesR, Fus, Nis pathways and the Lsr proteins of the Lux pathway. A more imited distribution across taxa is seen for the proteins of the two Com pathways and the Rcs and Agr pathways. Looking at each protein separately the most widespread proteins are CsrA, QseB, DesA, FusR, malt and LuxS (Fig. 2). Interestingly, those belong to different pathways (CsrA: UvrY, FusR: Fus, QseB: Qse, DesA: DesR, malt and LuxS: Lux pathway). The Lux and UvrY pathways contain the most widespread as well as some of the most limited protein types (LuxQ, Qrr and CsrB). There is also a difference in the distribution of the HKs and their cognate RRs: In most cases, the RRs are found in more bacterial groups than their cognate HKs (NarL, ComA, NisR, FusR, QseB, GlrR, and RpfG).

**Figure 2:**
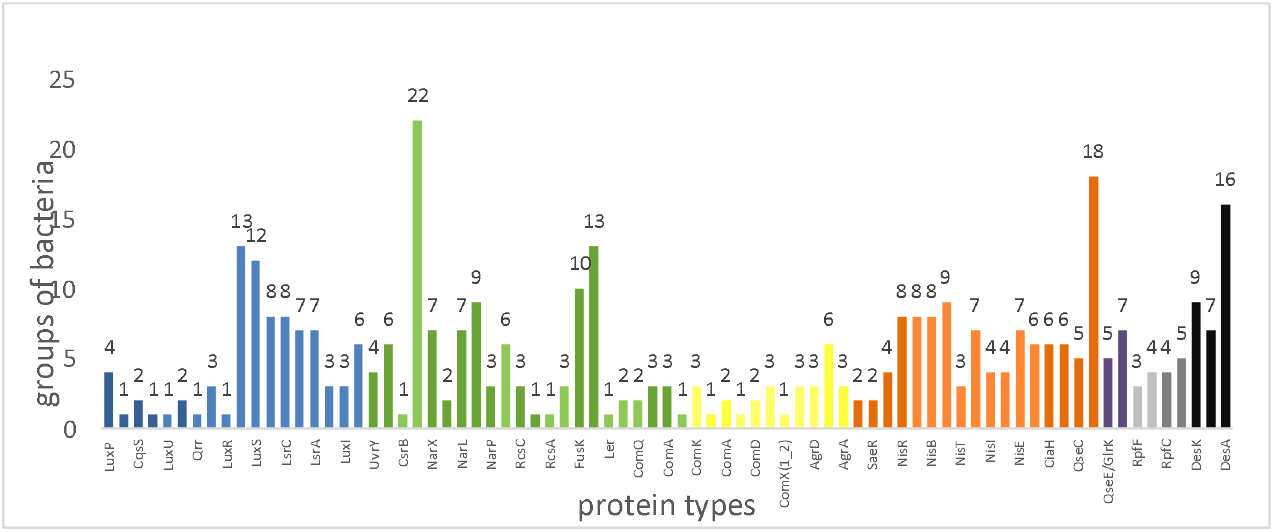
The number of bacterial groups in which each protein is found. The different colors indicate the family of each HK and RR in our study; proteins involved in the same pathway are indicated with a lighter shade of the same color. Blue: Lux family, Green: NarL family, Yellow: LytTR family, Orange: OmpR family, Purple: NtrC family, Grey: “Other”, Black: Des pathway.

### Conserved domains

In the Pfam database, the domains are organized into larger groups called “clans”. The NCBI database characterizes most domains as “superfamilies”. Table 1 contains the clans and their domains found in the studied amino acid sequences and Supplementary Table S2 shows the accessions for each domain in PFAM and InterPro, as well as the entry of the superfamily (NCBI-CDD) and pfam clan they belong to. The following domains do not belong to any clan: MASE1, PilJ, Hpt, Sigma54_AID, Trans_reg_C

**Table 1:**
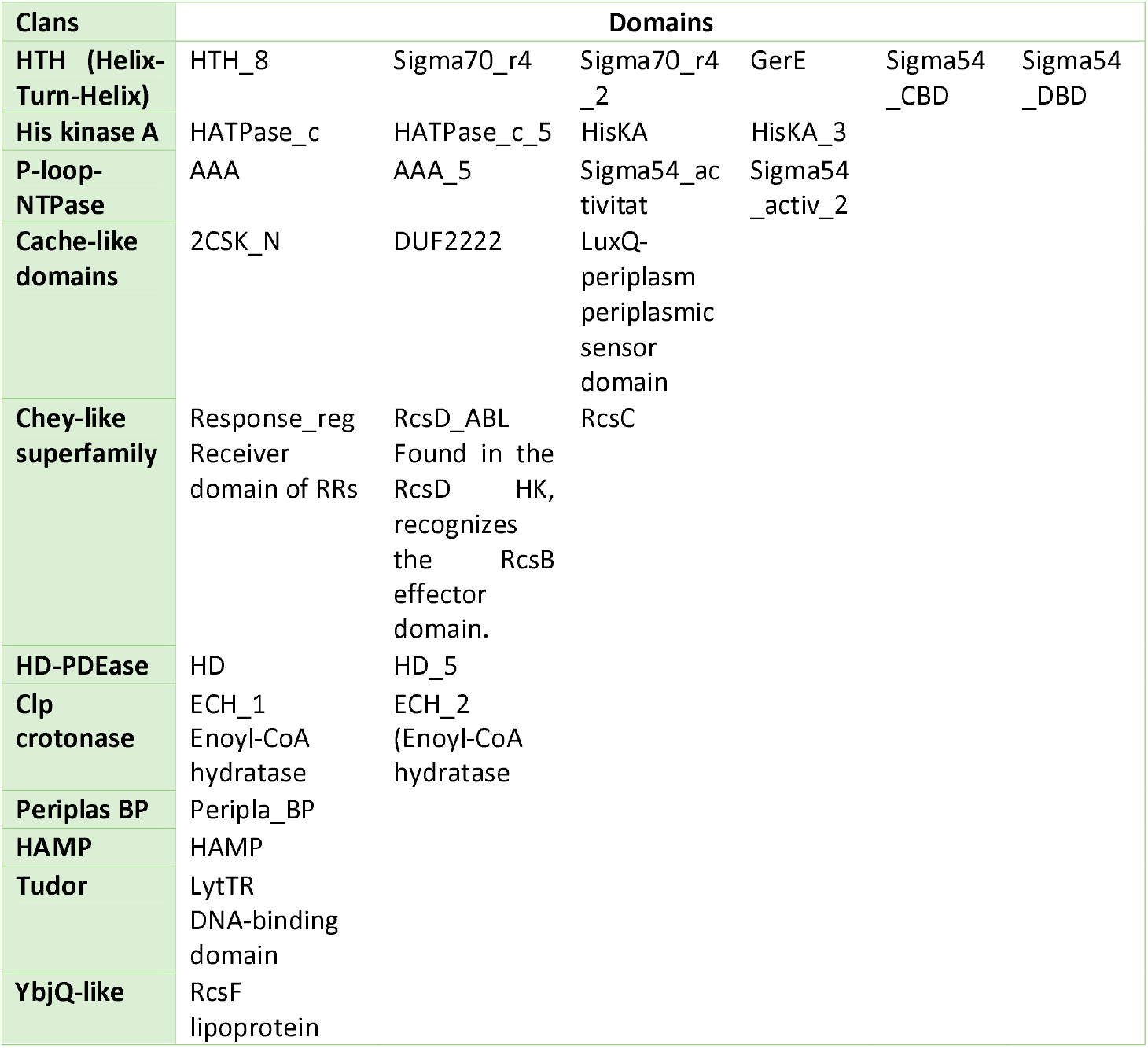
Categorization of the conserved domains of all the HK and RR types in this study.

#### Conserved domains in HKs

The length of the HKs varies from 406 (AgrC) to 949 (RcsC) amino acids. The longest HKs are the hybrid ones (NarL family: BarA, RcsC, Lux family: LuxN, LuxQ and CqsS, “Other”: RpfC) which also contain more conserved domains. Differences in the length of the sequences can also be found within the proteins of the same family. The HKs LuxQ and RcsC function as a complex with another protein (LuxP and RcsF, respectively).

Most HKs contain 1-3 conserved domains and up to 6 (Fig. 3). Comparing the number of the conserved domains, the members of the families OmprR, LytTR and NtrC (GlrK) have the lowest number of conserved domains (1 or 2). In contrast, the HKs of the NarL, Lux and “Other” (RfC) families have more than 2 conserved domains (Fig.3).

**Figure 3:**
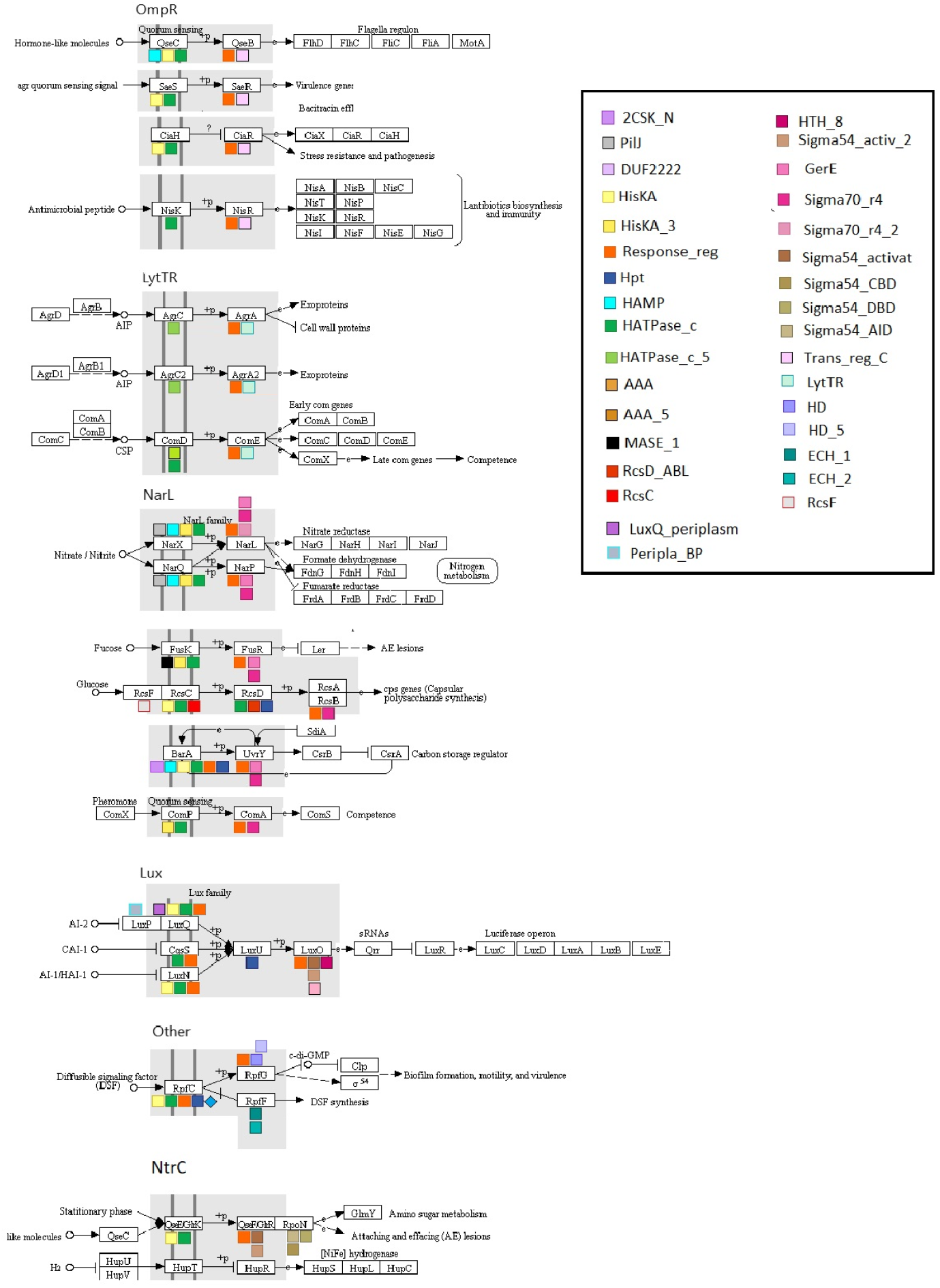
An overview of the position of the conserved domains in the HKs and RRs of this study. The similar colors of domains indicate their presence in a common clan of the Pfam database. The pathway drawings are from KEGG PATHWAY map ko02020, edited with Microsoft Draw.

The common trait is the existence of either HATPase_c or HATPase_c_5, which are also the only conserved domains in the protein NisK and the proteins of the LytTR family. According to NCBI-CDD, HATPase domains (Histidine kinase-like ATPase domains), also referred to as GHKL ATPase domains, are found in ATP-binding proteins, including histidine kinases. HAMP (Histidine kinase, Adenylyl cyclase, Methyl-accepting protein, and Phosphatase domain) is the second most frequent domain, found in HKs of the NarL family (NarX, NarQ, BarA). It relays various input signals but no common mechanism of HAMP has yet been established (NCBI-CDD). According to InterPro, one or more copies of the domain can be found in association with other domains, including HisKA. It has been suggested that its role is the regulation of the phosphorylation or methylation of homodimeric receptors by transmitting the conformational changes in periplasmic ligand-binding domains to cytoplasmic signalling kinase and methyl-acceptor domains. According to a study of Koretke et al., HAMP is the “linker region” and it is widely conserved in HKs as well as in chemoreceptors, bacterial nucleotidyl cyclases, and phosphatases. Its wide distribution is notable, as about a quarter of the HKs in the SMART database contain this domain and about half of all HAMP domains are found in HKs (Koretke et al., 2003). The Response_reg and Hpt domains are located in the C-terminal region of hybrid HKs, but only Hpt can alternatively be found as an independent protein (within the Rcs pathway). The Response_reg (Response regulator receiver domain or phosphoacceptor receiver-REC) receives the signal from the HK and Hpt (histidine-containing phosphotransfer domain), which mediates phosphotransfer reactions in Multi-step TCS signaling systems. Concerning the N-terminal region, only few proteins contain conserved domains, most of which belong to the NarL family (QseC, NarX, NarQ, FusK). These are Cache-like sensory domains (2CSK_N, DUF2222 and LuxQ-periplasm), PilJ (Type IV pili methyl-accepting chemotaxis transducer), Peripla_BP_4 (Periplasmic_Binding domain) and MASE1 (PFAM: predicted integral membrane sensory domain in HKs, diguanylate cyclases and other bacterial signaling proteins).

#### Conserved domains in RRs

The average length of RRs is significantly shorter that the HKs. It varies from 207 to 455 amino acids. Most RRs contain 2 conserved domains except for LuxO, which has 3 domains (Fig. 3). The N-terminal regions always contain the Response_reg domain, which is a common structure trait for all the RRs. The conserved domain on the C-terminal region is different in each RR family. The only exception is the protein GlrR (NtrC family), which has the same conserved domain as LuxO (Lux family). An overview of the conserved domains of both HKs and RRs is present in Table 2. HATPase_c is the most common conserved domain, as it is found in 15 protein types. HisKA is the second most common, with 9 occurrences.

**Table 2:** Occurence of the conserved domains of all the HKs and RRs in the different protein types and families.

| Domain | Protein Types | Family |
| --- | --- | --- |
| HATPase_c | 15 | All of the families |
| HATPase_c_5 | 3 | LytTR |
| HisKA | 9 | OmpR, NarL, Lux, Other, NtrC |
| HisKA_3 | 4 | NarL |
| HAMP | 3 | NarL |
| Response_reg | 3 | NarL, Lux, Other |
| Hpt | 2 | NarL, Other |
| Trans_reg_C | 4 | OmpR |
| LytTR | 3 | LytTR |
| GerE | 5 | NarL |
| Sigma70_r4_2 | 4 | NarL |
| Sigma54_activat | 2 | LuxO, NtrC |
| Sigma54_activ_2 | 2 | LuxO, NtrC |

Concerning the variety of the Effector domains, the majority has a DNA-binding activity: HTH_8, Sigma70_r4, Sigma70_r4_2, GerE, and Sigma54_DBD (DNA-Binding Domain) are HTH (Helix-Turn-helix) domains. Although Sigma54_CBD (Core Binding Domain) is also considered as an HTH domain, it interacts directly with the core RNA polymerase, forming an enhancer-dependent holoenzyme. The centre of this domain is slightly similar to a HTH motif, which may represent a DNA-binding domain. Also, some domains are P-loop-NTPases: Sigma54_activat, Sigma54_activ_2 interact with the sigma-54 factor of RNA polymerase; AAA and AAA_5 (ATPases Associated to a variety of cellular Activities) are described as a dynein-related subfamily and although their role is yet unclear, the AAA+ superfamily to which AAA ATPases belong, is associated with chaperone-like functions. Sigma_54_AID (Sigma_54_Activator interacting domain) is not classified into any clan. It is necessary for the interaction of Sigma54 RNAP holoenzyme with the activator and it can also inhibit transcription initiation prior to interaction with the activator.

### Phylogenetic trees by family

Given the families formed by the HKs and their cognate RRs, we constructed four phylogenetic trees for the HKs and three for the RRs. The HKs GlrK (NtrC family), RpfC (“Other”), their cognate RRs GlrR and RpfG, as well as LuxO are the only members of their family, hence no phylogenetic tree was inferred. Unless otherwise stated, only well-supported clades, with >90% bootstrap support, are discussed below. Phylogenetic reconstructions were done with PhyML using MAFFT alignments, but the major differences between the trees created from the MAFFT and MUSCLE alignments (Supplementary Fig. S6-S9) are also mentioned, wherever they apply.

#### The NarL family

The HKs of the NarL family are organized in distinct clades in the PhyML tree (Fig. 4A). Each clade consists of amino acid sequences of the same protein type, supported by high bootstrap values. The same structure is observed in the phylogenetic tree of their cognate RRs (Fig. 4B); the clades of the RR types are also supported by good bootstrap values, but most of them are below 95%. Although both trees have some amino acid sequences away from the clade of their protein type (e.g. marrComP, mdnBarA, paejNarQ and anxNarX), the phenomenon is more prevalent in the RRs. Notably, the structure of the phylogenetic tree shows three pairs of HK types (RcsC-BarA, ComP-FusK and NarQ-NarX), which group closer to each other than to the other protein types. This pattern is also seen for their cognate RRs.

**Figure 4:**
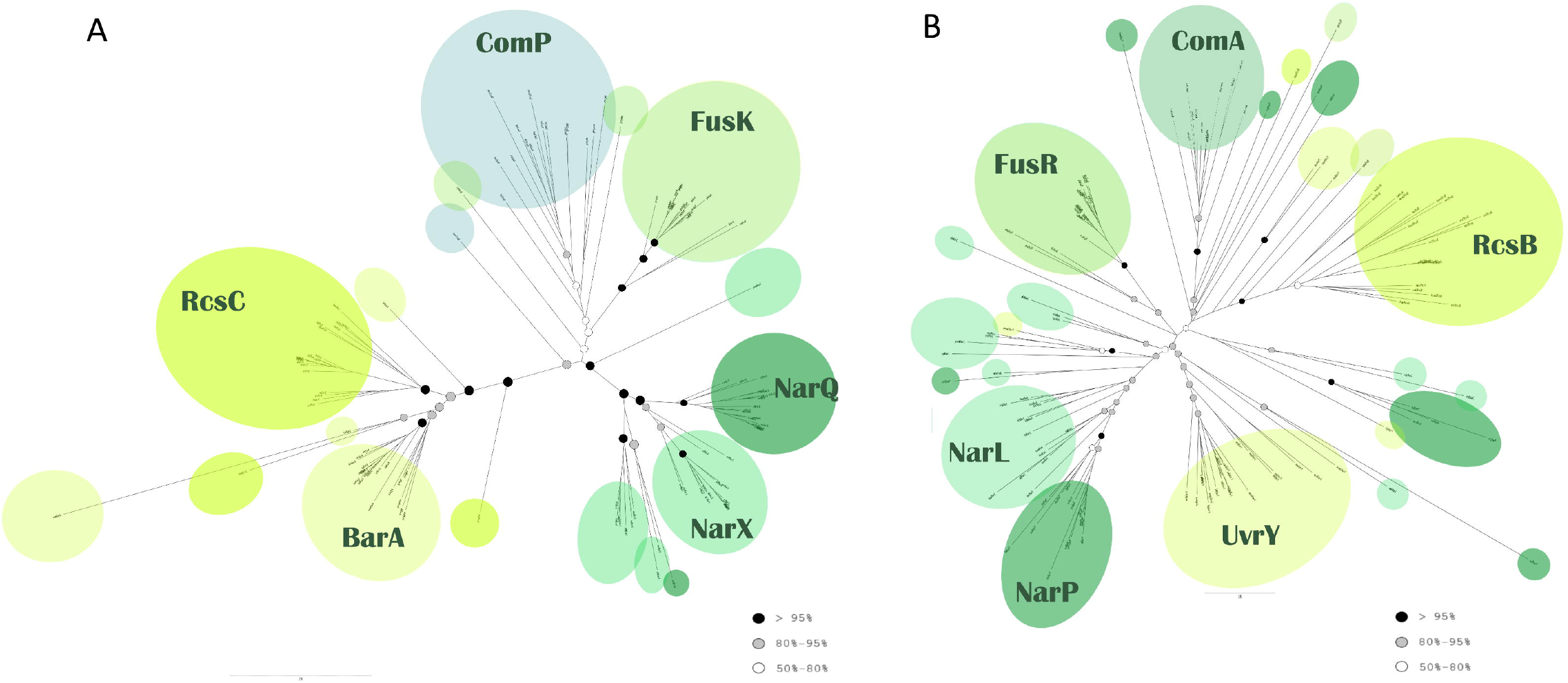
Phylogenetic analysis of the NarL family. (A) Histidine Kinases. (B) Response Regulators. Both trees reconstructed using PhyML based on the MAFFT alignment. The color of the bullets indicates the bootstrap value of the nodes. White: 50% -80%, Grey: 80%-95%, Black: over 95%

The clade of the RcsC proteins is closer to BarA than the rest of the protein types. On the other hand, their cognate RRs (RcsB and UvrY) are apart in the PhyML tree: although the UvrY proteins do not create a distinct pair with any other RR type, their clade is closer to FusR and the common clade of NarL and NarP. The RcsB RRs group together, except for the sequence abamRcsB. The UvrY proteins also form a clade (86% bootstrap support, excluding the first node of maesUvrY, nppNarL and ssilNarP sequences); outside of it sbxUvrY, dunUvrY and cbaeUvrY fall in the RcsB clade, mmarUvrY falls in the NarL clade, and tjeUvrY in the NarP clade. A comparison with the same PhyML tree from the MUSCLE alignment (Supplementary Fig. S6B) shows that although the distribution of the NarL and NarP RRs is in multiple dispersed clades in the tree, the UvrY and RcsB proteins clearly form a common clade (87% bootstrap value). Also, the ComA clade is closer to FusR than to RcsB.

The ComP clade has a 50% bootstrap value, excluding kbsComP and marrComP, and it groups with the FusK clade (72% bootstrap value). In contrast, the clades of their cognate RRs, ComA and FusR, do not group together. The ComA sequences are not directly related to any other clade but they are close to RcsB in the tree.

The NarX proteins are separated in two main clades, both of which share a common node with the NarQ clade; spswNarX branches out earlier. However, the distribution of their cognate RRs NarL and NarP is more complicated. They are divided into several clades (sometimes including both protein types), yet most of these clades are placed close to each other and their common node has a 79% bootstrap value. In detail, the NarP sequences are grouped into two distant main clades; a larger one, neighboring with the NarLs and a smaller distant one. There are more sequences outside of these clades than is seen for their cognate HK sequences, and from different species than in the HKs tree. The NarLs are gathered in several small clades close to each other, most of which demonstrate bootstrap values of at least 80%, but many NarL sequences are dispersed throughout the phylogenetic tree.

#### The OmpR family

All different HK types of the OmpR family are grouped into separate clades. None of the HK sequences is misplaced in a clade of a different protein type, except spyaNisK, which falls in the CiaH clade. Their cognate RRs follow a similar pattern, except for sequences phmQseB and aprsQseB, which lie outside the QseB clade. There are two related groups in both phylogenetic trees: CiaH -QseC and NisK – SaeS and their cognate RRs, CiaR -QseB and NisR – SaeR (Fig. 5A and 5B). The majority of the CiaH sequences stem from a common node, with soxCiaH and cooCiaH branching out earlier. The QseC sequences are distinctly separated from the rest of the RR types and they share a common node with the CiaH clade. The cognate RRs follow the same pattern: The clade of the QseB sequences (85% bootstrap value) forms a common clade with the CiaR sequences. The NisK sequences are organized into one main clade and they have a common node with all the SaeS proteins. The NisR proteins also group together with the clade of the SaeR sequences.

**Figure 5:**
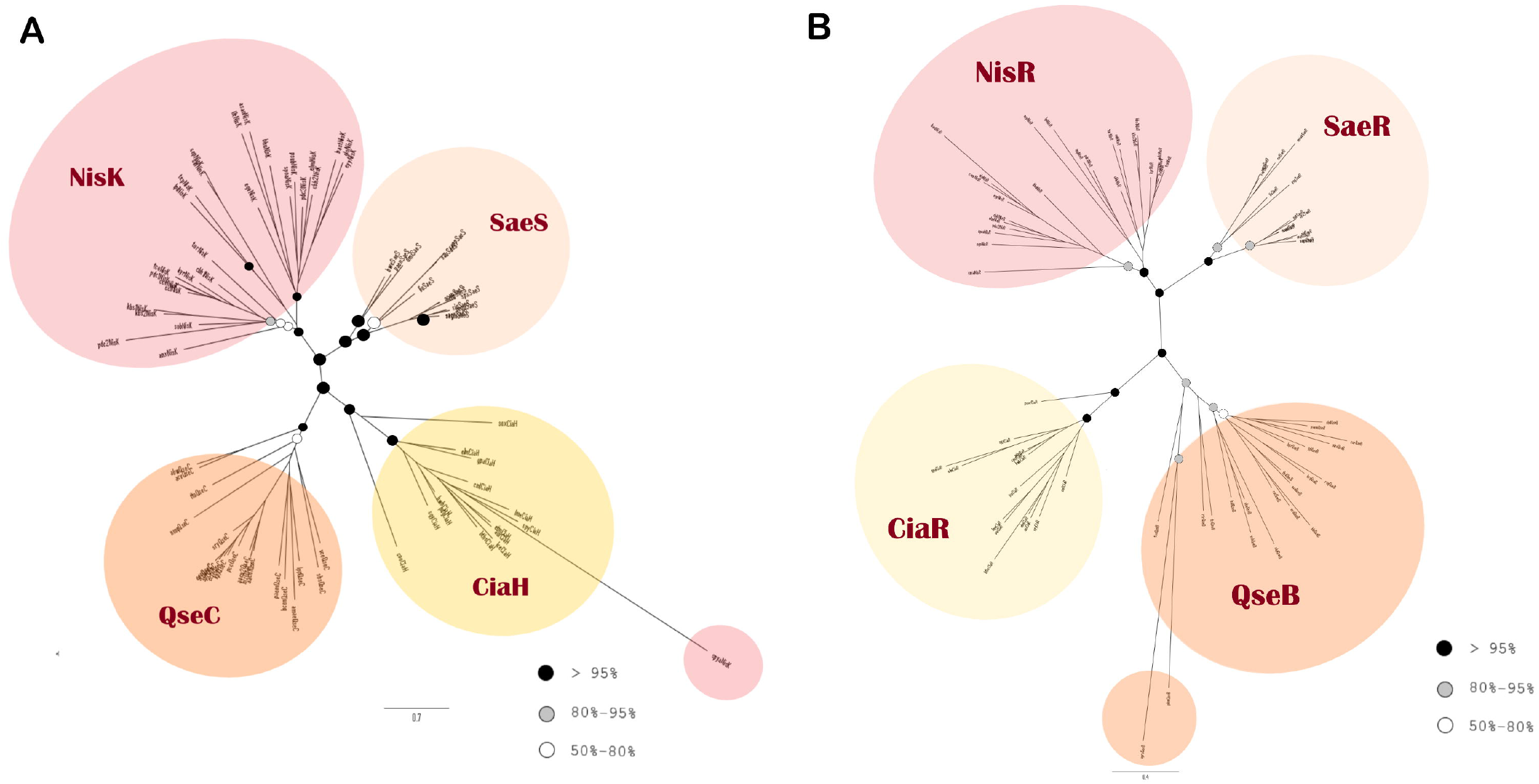
Phylogenetic analysis of the OmpR family. (A) Histidine Kinases. (B) Response Regulators. Both trees reconstructed using PhyML based on the MAFFT alignment. The color of the bullets indicates the bootstrap value of the nodes, as in Figure 4.

#### The Lux family HKs

The sequences of each HK type of the Lux family (LuxN, LuxQ and CqsS) are grouped into distinct clades (Fig. 6). The LuxN sequences have one common node. On the other side of the tree, LuxQ create a common clade, although dhyLuxQ, dsaLuxQ, oceLuxQ are more diverged. The CqsS sequences branch out at the center of the tree in 3 clades, with fsoCqsS more diverged. On the contrary, most LuxQ branches in the PhyML tree of MUSCLE (Supplementary Fig. S8) appear near the core of the tree; it seems that the clades CqsS and LuxQ switch positions in the MAFFT and MUSCLE versions of the tree. As there is only one RR type for all the Lux family HKs (LuxO), there is no phylogenetic tree for the RRs of the Lux family.

**Figure 6:**
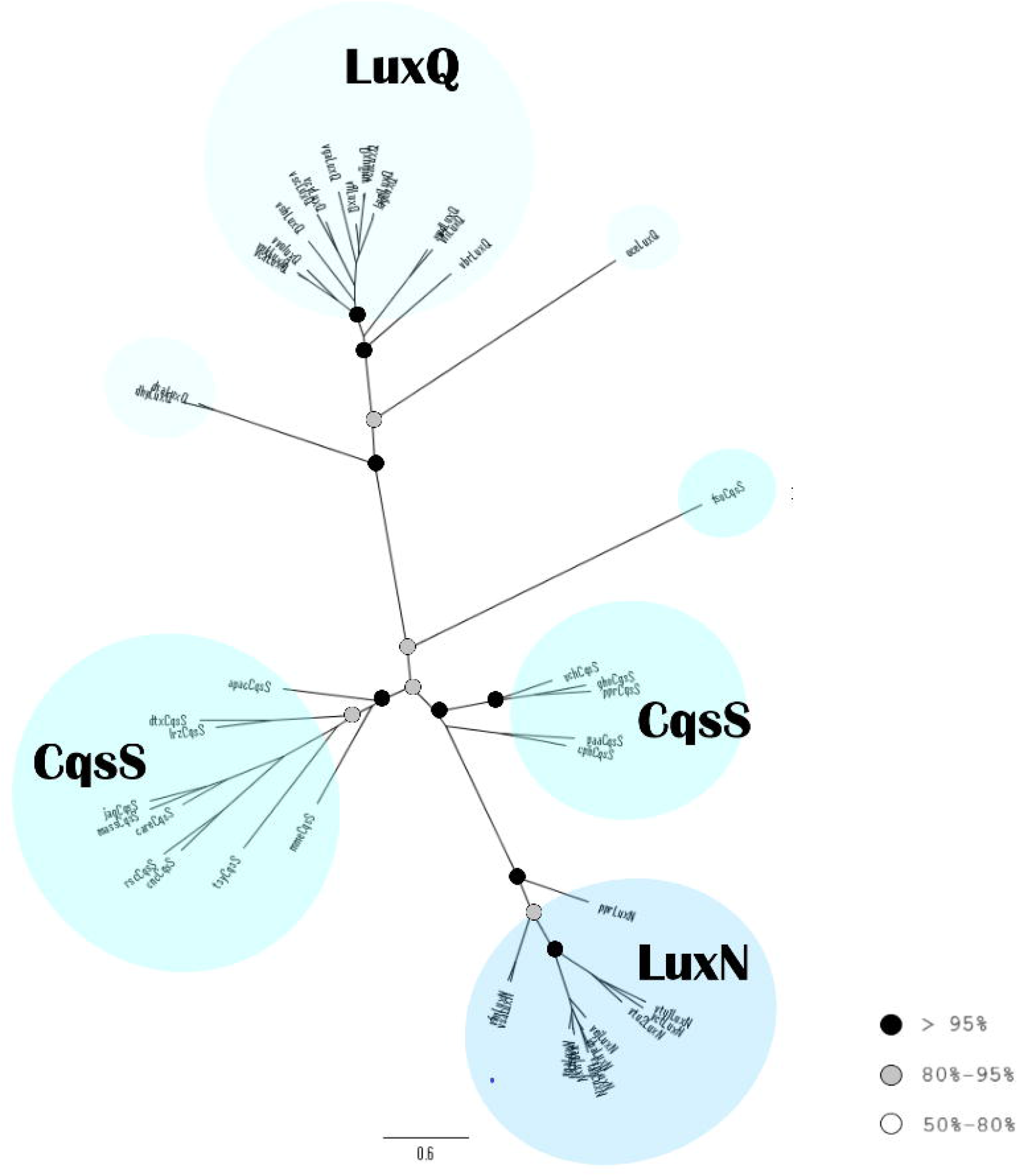
Phylogenetic analysis of the Lux family histidine kinases. Tree reconstructed using PhyML based on the MAFFT alignment. The color of the bullets indicates the bootstrap value of the nodes, as in Figure 4.

#### The LytTR family

The different types of HKs of the LytTR family, as well as their cognate RRs, are clearly separated from each other and both phylogenetic trees have a similar overall appearance (Fig. 7A and 7B). The AgrC sequences form several clades, all of which stem from a common node on the phylogenetic tree. The two AgrC2 sequences are found in a distinct clade within the AgrC clade, closest to rimAgrC. Similar to AgrC HKs, the AgrA sequences do not form one distinct clade, but smaller ones, gathered on one side of the phylogenetic tree. The AgrA2 sequences are found in a distinct clade near the AgrA clades closest to cdfAgrA (77% bootstrap value). Most of the ComD sequences are gathered in one clade, apart from the lhiComD, lsjComD and laliComD sequences, which are placed among AgrC sequences. The ComE RRs create a main common clade, with the exception of an earlier smaller clade: sequences std2ComE and smb1ComE branch early (58% bootstrap support) as do their cognate HKs.

**Figure 7:**
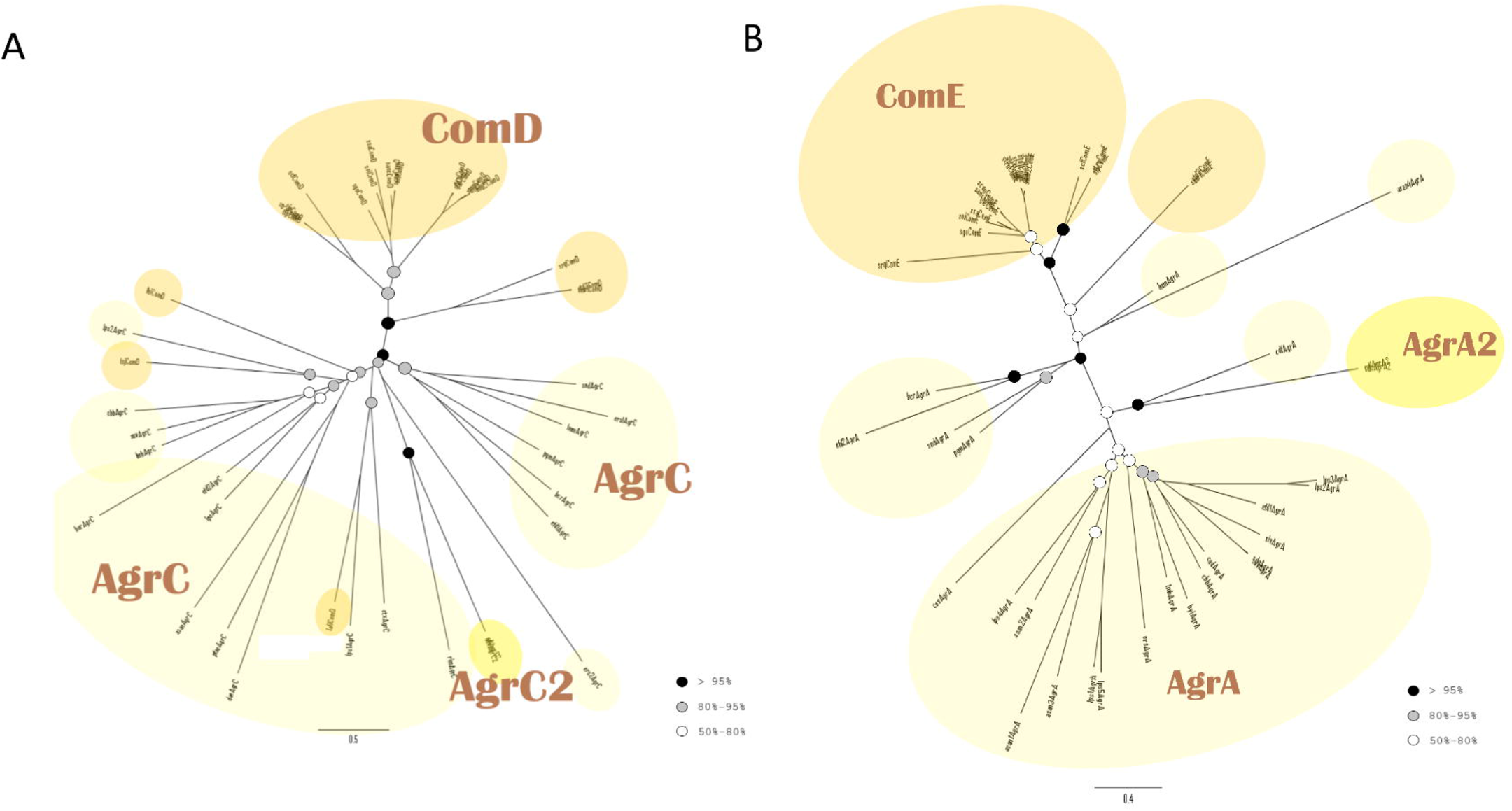
Phylogenetic analysis of the LytR family. (A) Histidine Kinases. (B) Response Regulators. Both trees reconstructed using PhyML based on the MAFFT alignment. The color of the bullets indicates the bootstrap value of the nodes, as in Figure 4.

### Phylogenetic trees of all the families together

#### Phylogenetic analysis of all the HK families

The two phylogenetic trees of all the HKs and RRs families together are discussed below, based on the MAFFT alignment. Major differences with the phylogenetic trees based on the MUSCLE alignments are also mentioned. In general, the members of each HK family tend to form one main clade, although the bootstrap values are not always significant. The clades of each HK protein type are also mostly monophyletic (Fig. 8). The sequences of the LytTR family, as well as the RpfC (“Other”) and GlrK HK types (NtrC family) are each gathered in one clade, whereas the protein sequences of the Lux as well as the NarL family are found in more than one clades. The following 3 groups of families are discernible: 1) LytTR and NarL (NarQ, NarX, FusK and ComP), 2) OmpR and NtrC, 3) Lux, NarL (BarA, RcsC) and “Other” (RpfC).

**Figure 8:**
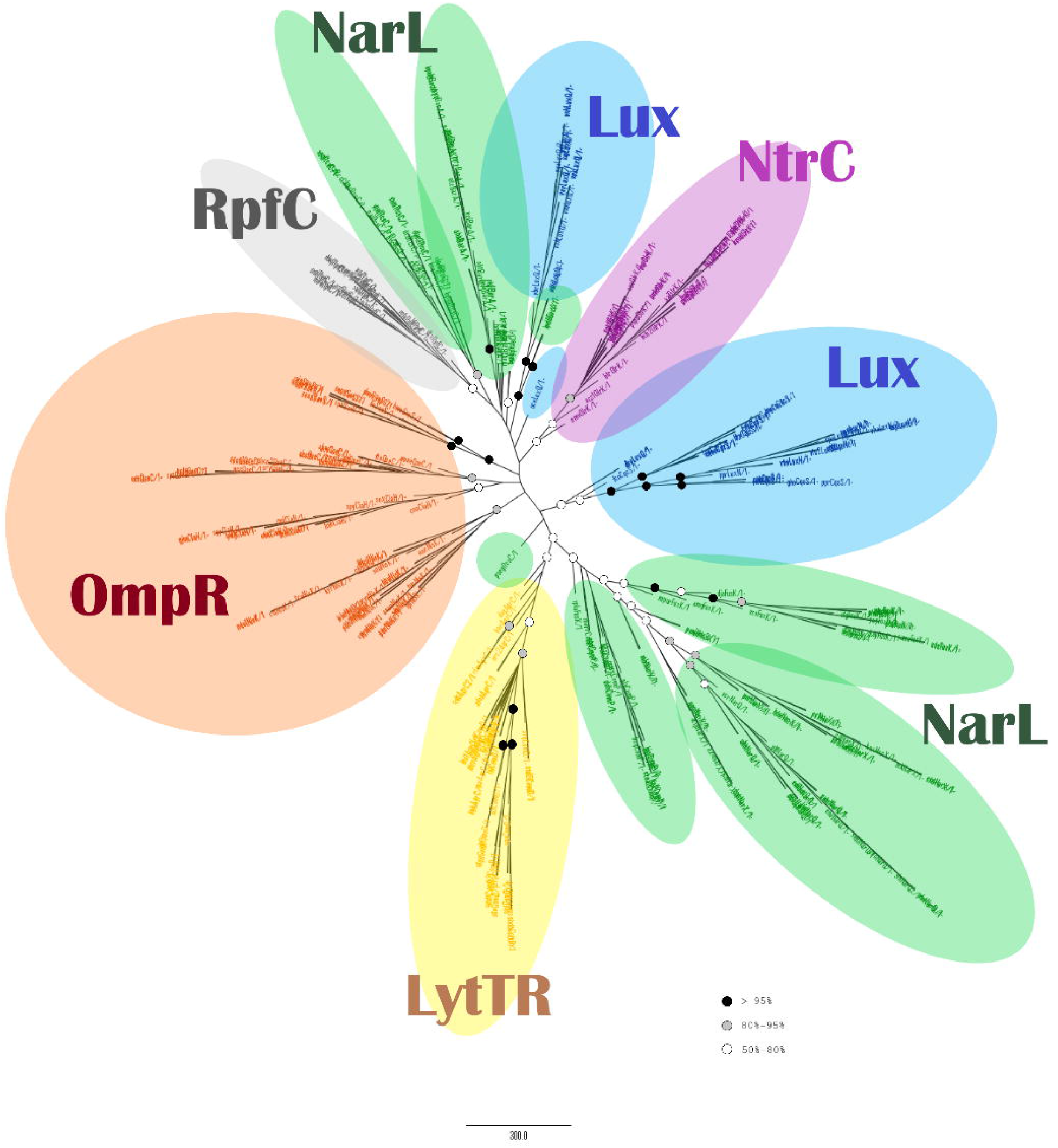
Phylogenetic analysis of all histidine kinase families. Tree reconstructed using RaxML based on the MAFFT alignment. The bullets indicate the bootstrap value of the nodes, as in Figure 4.

##### NarL family

The sequences of this family form three main clades: the RcsC clade, the BarA clade (97% bootstrap value excluding the early branches) and a third -and larger-one which includes the NarQ, NarX, FusK and ComP sequences (Fig. 8). Within this larger group, the FusK sequences cluster together (56%) and create a common clade with the NarQ and NarP clades. All the ComP sequences also derive from a common node, but with very low bootstrap value. NarQ sequences form a well-supported main clade, while NarX forms two main clades with 87% and 97% bootstrap support, respectively. Only one NarQ and three NarX sequences fall outside their protein type clades. In general, the grouping of the clades of the NarL family is very similar to the tree for the HKs of the NarL family alone (Fig. 4): the BarA-RcsC, and NarX-NarQ pairs are recapitulated, and the only difference is the positions of FusK, which is closer to ComP in Fig. 4, while it shares a common node with the NarQ/NarX clade in Fig. 8.

##### OmpR family

The HK types of this family are separated into three main clades: one for QseC and CiaH one clade for the SaeS and one for the NisK sequences (Fig. 8). The tree from the MUSCLE alignment (Supplementary Fig. S10) recapitulates the exact structure of the phylogenetic tree of the OmpR family alone (Fig. 5), as SaeS and NisK sequences create a common clade. The clades of each protein type of the OmpR family are supported by high bootstrap values, although their common node with the rest of the phylogenetic tree has low bootstrap support.

##### Lux family

Two distinct clades and a smaller mixed one are formed by the HKs of the Lux family: one clade consists of CqsS and LuxN sequences (with dhyLuxQ and dsaLuxQ at the root of the clade) and neighbors with the OmpR, LytR and NarL clades and the other is made of only LuxQ sequences near the BarA clade of the NarL family (Fig. 8). This pattern is also seen in the tree based on the MUSCLE (Supplementary Fig. S10), and follows the groupings of the phylogenetic tree of the Lux family alone (Fig. 6), as CqsS and LuxN are grouped together, while the LuxQ sequences are placed away from the rest of the members of the Lux family.

##### LytTR family

All the sequences of this family derive from a common node (56% bootstrap support) (Fig. 8). The ComD sequences form one clade (88%) within the AgrC clade. The AgrC sequences form many small clades, including the two AgrC2 sequences. The closest clade to the LytTR family is the overall clade of the ComP-NarX-NarQ-FusK sequences of the NarL family (62% bootstrap support).

##### NtrC family (GlrK proteins-QseE)

All the sequences of the GlrK (QseE) HK are in one clade, with 73% bootstrap support (Fig. 8). The closest clade to them is that of SaeS (of the OmpR family), but with insignificant bootstrap support; in the tree based on the MUSCLE alignment, the NtrC family groups more clearly with the OmpR family, and in particular the common clade of QseC and CiaH, although again with very low boostrap support.

##### “Other” (RpfC sequence)

All RpfC sequences are placed in one clade, with 74% bootstrap value. The neighboring clade consists of RcsC sequences of the NarL family (Fig. 8), but the bootstrap value for their common clade is very low.

#### Phylogenetic analysis of all the RR families

The RR families are distinctly separated from each other in the phylogenetic tree. Also, it is clear that the sequences of each RR type within each family remain together, like in the phylogenetic tree of each family. Very few sequences fall outside of the clade of their family or RR type (Fig. 9). An overview of the tree reveals a closer relationship between the following family groups: 1) the RpfG and NarL families, 2) the OmpR, NtrC and Lux families and 3) the LytTR clade, which is closer to group 1). In the tree based on the MUSCLE alignment (Supplementary Fig. S11), the position of the RpfG (Other) group is closer to NtrC and Lux families; however, the position of this group relative to other families is not supported by high bootstraps in either of the trees.

**Figure 9:**
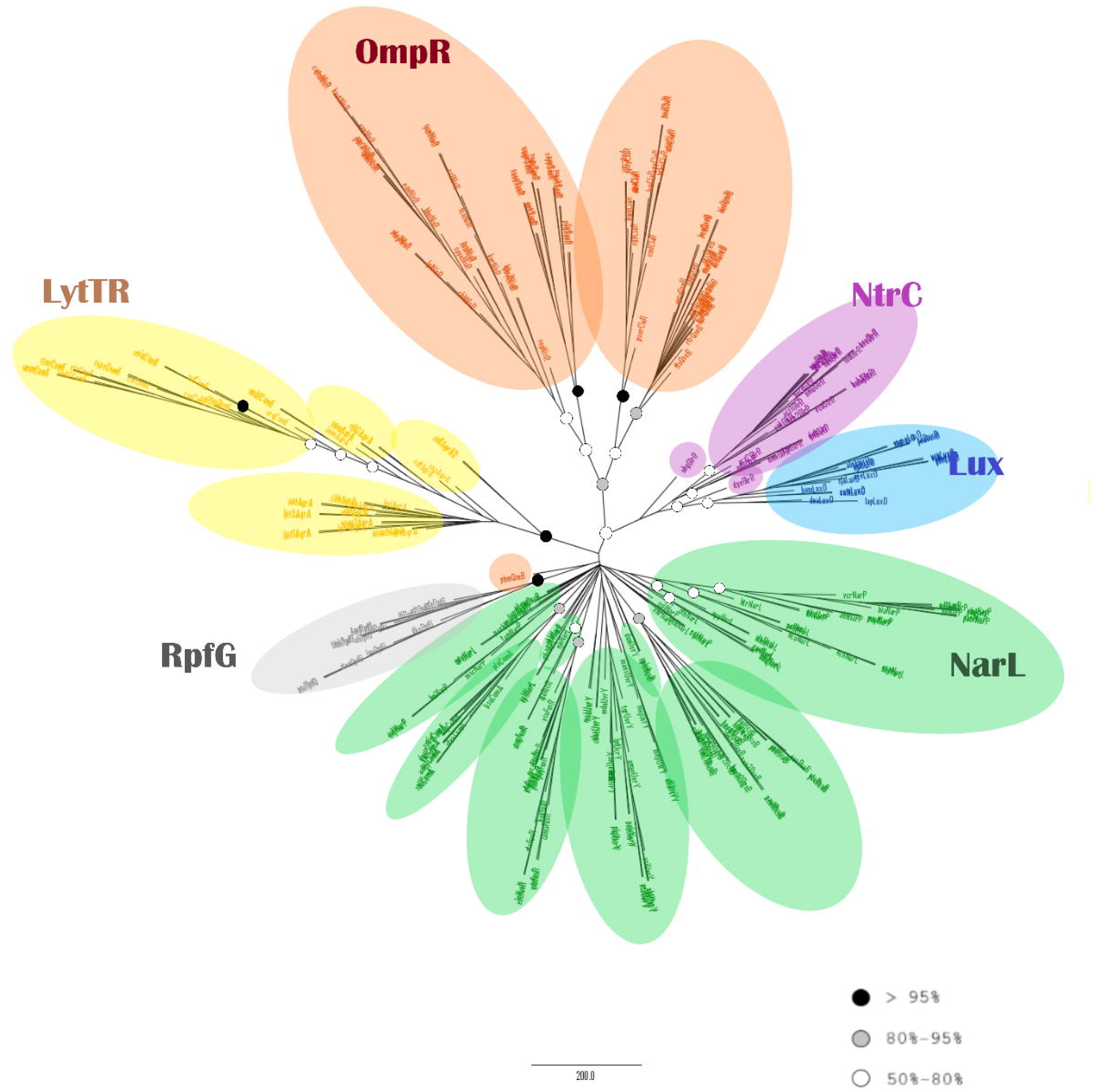
Phylogenetic analysis of all response regulator families. Tree reconstructed using RaxML based on the MAFFT alignment. The bullets indicate the bootstrap value of the nodes, as in Figure 4.

##### NarL family

The NarL family members create a sum of clades, each consisting mostly of sequences of one particular NarL protein type. The positions of all the family members are very close to each other, to the extent that there are no defined pairs of protein types. The RcsB and ComA clades branch later in the NarL family. All RcsB and ComA RRs are grouped in their own RR type clade, with the exception of abamRcsB. The clade of the UvrY sequences has, in general, nodes of low bootstrap value and five sequences are placed outside of it: tjeUvrY, dunUvrY and mmarUvrY. All the FusR RRs form one clade, except spirFusR and sphnFusR, which are right before the clade of the RcsB proteins. Most NarPs are divided into two main clades, one with NarL sequences and a second one near the ComA sequences. The NarL sequences are widely distributed in the clades of the NarL family, forming several small clades. One group of these clades is close to the NarP sequences and another group is placed among the ComA and FusR clades.

##### OmpR family

The members of the OmpR family are grouped together (86% bootstrap value) and there is one individual clade for each protein type, supported by significant bootstrap values (except NisR: 63%). SaeR and NisR group together with 72% bootstrap value and CiaR with QseB (58% bootstrap). This recapitulates the pattern seen in Fig 5B. Only phmQseB is distant in the tree, near th RpfG clade (but with inisgnficant bootstrap support).

##### Lux family

The Lux family (LuxO) (78%) is close to NtrC, forming a common clade (52%). This clade shares a common node with the OmpR family (69%).

##### LytTR family

The clade for all the RRs of the LytTR family is closer to the NarL and RpfG family members and it consists of many smaller clades or individual branches. The different RR types belong to separate clades where the AgrA2 sequences branch out earlier of all and they have a common node with the AgrA clade (100%), wheras the ComE is the last clade.

##### NtrC family (GlrR protein)

The GlrR sequences group together; however, the early nodes have a very low bootstrap value. The obgGlrR sequence branches out early with a very low bootstrap value (32%), while dyeGlrR lies outside the NtrC clade, on the LuxO clade, with 78% bootstrap value.

##### “Other” (RpfG protein)

The RpfG RRs form one clade with 96% bootstrap value, which includes no sequences of other RR type. The closest RR family is NarL, complying with the structure of the tree of all the HKs (Fig. 8).

## Discussion

The QS mechanism includes a multitude of signal molecules and pathways, which aim to coordinate the activities of a bacterial population by changing gene expression according to the population density. The fact that this mechanism affects various processes makes this mechanism a potential target in antibiotics research. Few phylogenetic analyses on the bacterial QS exist, and given the multiple pathways contributing to QS, studies have so far focused only on τhe signaling molecules, or on specific pathways. Among them, the LuxI/R pathway (Gram positive) has been mainly studied. Based on the lack of a wider phylogenetic study of QS proteins, we chose to analyze proteins of the QS pathways which belong to a widespread signal transduction system, the Two Component System (TCS). This mechanism includes pairs of a Histidine Kinase (HK) and its cognate Response Regulator (RR). The studied QS pathways are found in a wide variety of both Gram-positive and Gram-negative bacteria.

### QS protein conservation

Looking at the distribution of the different QS pathways across bacterial taxa, the overall conclusion is that α-, β-, γ-proteobacteria, clostridia and bacilli possess the highest number of the QS proteins (Fig. 1). The most widespread proteins are CsrA, FusR, QseB, DesA, malt and LuxS. These belong to various pathways and families and they have different roles, indicating no clear relationship between these characteristics and the number of the bacterial groups containing them. The least widespread proteins are found in the two Com pathways, the Rcs and Agr pathways, as well as the Lux pathway. These results suggest that the wide presence of one protein doesn’t mean a similar distribution for the rest of the proteins of the same pathway. This contradiction is pointed out by the fact that the Lux and UvrY pathways contain both the most widespread and some of the most limited protein types (LuxQ, Qrr and CsrB). Also, there is a difference even in the distribution of the HK/RR pairs, i.e. between the HK and its cognate RR. This distribution may simply be the result of inadequacies in genome annotation, or it may be an indication of flexibility in the system, meaning that HKs can function with other than their “cognate” RRs, in essence leading to mixed pathways. Although TCS pathways demonstrate a high specificity overall, such “cross-talk” between different TCS pathways is possible (Agrawal et al., 2016). In the case of “orphan” HKs or RRs with no cognate protein (Freedman et al., 2019; Wang et al., 2009), an interaction between an orphan protein with a non-cognate one can also occur (Wang et al., 2009). Also, given the QS map of KEGG (Fig. 3), QseC can interact with GlrR (QseF), which is not only a non-cognate RR, but it also belongs to a different family (NtrC); interestingly, QseC and GlrK both bind to Adrenaline and Noradrenaline. All of the interactions depicted in the QS maps of KEGG (maps ko02020 and ko02024) involve proteins of the OmpR family. QseB inhibits the cognate pair of FusK/FusR of the NarL family, CiaR inhibits ComC, ComD and ComE proteins of the LytTR family and SaeS is activated by the agr QS signal.

Examining each protein family of our study separately, it seems that the most widespread HK/RR pairs of QS are those of the NarL, OmpR and NtrC families, whereas the Lux and LytTR are the least common, although the Lux pathway is the most well-known and well-studied QS pathway.

### Conserved Domains

The HK-RR pairs belong to 6 families, as defined by KEGG: OmpR, LytTR, NarL, Lux, NtrC and “Other”. In both the OmpR and the LytTR family, the different protein types consist of a small number of common conserved domains. On the other hand, HKs of the NarL and Lux families show common domains and a number of variable domains; notably, the NarL and Lux families also include hybrid HKs. The RRs of all families contain common domains (all of them have the Helix-Turn-Helix motif).

The Lux family consists exclusively of Hybrid HKs, and all 3 of them activate a common phosphotransfer protein and RR. The NtrC contains only the GlrK HK, which has the same conserved domains with the HKs of the OmpR family, a characteristic which is also evident in their close relationship on the phylogenetic tree. However, its cognate RR (GlrK) has various common domains with the RR of the Lux family. This is also evident in the phylogenetic trees. Therefore, we conclude that similarities can be found not only among the protein types within a family, but also among different family members, which indicate a possible common evolutionary history.

There is partial agreement of our results with previous sudies on the classification of HKs and RRs. The differences are mostly due to differences in the selection of proteins included in each study, and due to differences in classification based on domain structure alone versus phylogenetic analysis. For example, looking at the classification of the HKs in the Grebe and Stock study, ComP, NarQ and NarX were classified in group HPK7, CiaH in HPK1a, NisK in HPK3c and BarA, RcsC, RpfC in HPK1b (Grebe and Stock, 1999). However, only some of the HKs mentioned here were also included in that study, and this way of classification differs from the KEGG database and our phylogenetic trees results, as CiaH and NisK (OmpR family members) belong to separate groups and RpfC (hybrid HK) is placed with the hybrid HKs of the NarL family (although hybrid HKs were placed in other HPK groups as well).

In the study of Forst and Kim, BarA and RcsC were characterized as Type IB and NarX and NarQ as Type III HKs (Forst and Kim, 2015). However, no members of other families included in our study were present in that analysis, so we cannot conclude if the comparison criteria used by Forst and Kim led to a different classification or if their classification is more detailed and separated the members of one family into multiple groups. Capra and Laub classified the HKs into 4 groups depending on the presence of PAS, HAMP and GAF domains or the absence of these domains (Capra and Laub, 2012). According to the authors, most HKs have at least one domain between the transmembrane part of the protein and the Histidine phosphotransfer domain, with PAS, HAMP and GAF being “by far the most common” ones. However, our studied proteins had no PAS and GAF domains, with the exception of RcsC, in which the PAS domain had a very high E-value in KEGG and SMART and thus it didn’t meet our criterion to be included in the conserved domains of our study. Nevertheless, all three HKs with a HAMP domain (NarX, NarQ, BarA) were grouped to the same family (NarL), in agreement with the Capra and laub study. What contradicts their classification is the fact that the NarL family contains both HKs with HAMP and others without a HAMP domain, which are supposed to belong to different groups according to Capra and Laub. Similarly, the effector domains for the classification of the RRs were also different in the research of Capra and Laub, as no GGDEF and Methyltransfer domains were present in the proteins of our study. However, the LuxO and GlrR proteins possessing the AAA_5 domain, were grouped close to each other in our phylogenetic trees, complying with the classification criterion of Capra and Laub.

Concerning the classification of the RRs according to Hakenbeck and Stock (Hakenbeck and Stock, 1996) it is the same as the data of KEGG and our study. CiaR and NisR were classified to the OmpR family and ComA, NarP, NarL and RcsB were classified to the FixJ family. The results of Grebe and Stock are similar(Grebe and Stock, 1999): CiaR and NisR were classified to the RA1 group and ComA and RcsB to RE, but NarP and NarL were classified to a different group (RC1). In this paper, AgrA and ComE are also mentioned and grouped to RD, both of which belong to the LytTR family in our study.

### Phylogenetic trees

In all the inferred phylogenetic trees of each family, the amino acid sequences of both HKs and RRs are grouped in separate distinct clades according to their protein type and in many cases, with high bootstrap value. The same happens in the phylogenetic trees of all families together, with distinct clades for sequences of different family and protein types. Also, the relation among protein types (either HKs or RRs) is generally the same in the tree of each family and of all families together. In most cases, the relations among the HKs are also reflected by the relations of their cognate RRs, indicating a common evolutionary history, as has already been suggested by former studies on other proteins of the TCS (Koretke et al., 2000).

The members of the NarL family form pairs of HKs that are closely related, NarX-NarQ, BarA-RcsC and ComP-FusK in both the tree of the NarL family alone and the tree of all the HK families together. In both trees the BarA-RcsC group is more distant. The tree for all the families together placed all RR types of the NarL family very close to each other: NarL and NarP clades are clustered together and UvrY sequences are placed close to RcsB, being in agreement with the trees of their cognate HKs. However, in the RR tree of the NarL family alone, only NarL and NarP demonstrate a similar grouping to the HKs from which they are activated: ComA is closer to RcsB (instead of FusR), and UvrY is closer to the clade of NarL/NarP (instead of RcsB).

The different protein types of the OmpR family demonstrate the same relation in all the phylogenetic trees (of the OmpR family alone and the tree with all the HK families together). There are two pairs of HKs: CiaH-QseC and SaeS-NisK. Their cognate RRs follow the same relation in all the RRs phylogenetic trees: the SaeR clade is closer to NisR and the CiaR clade is closer to QseB. Regarding the Lux HKs, both the tree of the Lux family and the tree for all the HKs show a similarity between the CqsS and LuxN. The latter also places the LuxQ sequences in the clade of other families (BarA, RpfC and RcsC sequences). Their common RR (LuxO) is in the same clade with GlrR in the tree of all RRs. The protein members of the LytTR family have separate clades for the AgrC -ComD and ArgA-ComE sequences in all trees. The AgrC2 and AgrA2 sequences belong to the AgrC and AgrA clade, which is confirmed by all the inferred phylogenetic trees. The position of the RpfC/RpfG and GlrK/GlrR sequences (NtrC family) are slighlty different for the HKs and their RRs. The RpfC sequences are closer to RcsC, whereas RpfG is closer to NarL and NarP. GlrK is closer to the OmpR family but GlrR is in the same clade with LuxO.

A comparison of the phylogenetic trees of all HKs and all RRs suggests two clusters of the QS protein families: 1) OmpR-NtrC-Lux,and 2) NarL and LytTR, as well as clusters of protein types per family as outlined above. These evolutionary relationships highlight a common evolutionary history, and can inform future applications, such as the design of novel inhibitors for pathogenic QS systems. In fact, there have already been numerous scientific papers about Quorum Quenching, including for some of the proteins of our study. Diol-containing compounds, boronic acids and sulfones have been suggested as antagonists of the signaling molecules which bind with LuxP and as potential solutions to biofilm formation (Brackman and Coenye, 2014). Other molecules inhibiting the QS pathways of Gram-negative pathogens have been reported, including for the HKs and RRs of the Lux family and RpfC/RpfG, by blocking the receptors, inhibiting the production of the signaling molecules, or by degrading them (Defoirdt, 2018). The inhibition of the Agr pathway of *Staphylococcus aureus* has also been studied, with natural and synthetic substances targeting either AgrC or AgrA: the cyclodepsipeptide Solonamide B of *Photobacterium halotolerans* interferes with the binding of AIPs to AgrC, whereas ω-hydroxyanodin from *Penicillium restrictum* binds to AgrA and therefore inhibits the expression of the regulated genes (Scoffone et al., 2019). Savirin (*S. aureus virulence inhibitor*) is another small molecule that selectively targets AgrA (Sully et al., 2014). The synthetic small molecule inhibitor LED209 is selective to QseC (Curtis et al., 2014). The genes ComD (HK) and ComE (RR) was also found to be downregulated by a small molecule containing a 2-aminoimidazole subunit (2A4) along with five other biofilm-related genes of *S. mutans* (Liu et al., 2011). The antiobiotic mupirocin when used in sub-inhibitory concentrations on high-level mupirocin-resistant MRSA strains (Methicillin-resistant Staphylococcus) interferes with the agrA and saeRS genes (Jin et al., 2018). Our prediction is that such inhibitors are more likely to also interfere with protein types similar to the original target protein, as highlighted by the phylogeneti analysis, than with more distant ones, e.g. an inhibitor for a Lux family HK may also affect NtrC family HKs. On the other hand, pathway cross-talk in species with multiple QS systems may hinder these inhibitory effects. Broader inhibition approaches have also been suggested, with the ATP-binding domain of HKs (HATPase domain) being the main target for TCS pathways inhibition (Wilke et al., 2015; Worthington et al., 2013).

Two-component signal transduction arose early in bacterial evolution after their separation from the last common ancestor with significant diversification throughout the subsequent bacterial speciation; HKs arose from ATPases of the GHKL superfamily and pyruvate dehydrogenase kinases (PDKs), while the origin of the RRs remains unclear (Capra and Laub, 2012; Koretke et al., 2000). Regarding the origin of two-component quorum sensing systems, a phylogenetic analysis for the proteins LuxI, LuxR and LuxS in firmicutes, proteobacteria and actinobacteria, indicated that these proteins appeared very early in the evolutionary path of the bacteria, but with multiple occassions of horizontal gene transfer resulting in their present-day distribution (Lerat and Moran, 2004). In a study of the ComQXPA pathway in firmicutes, the phylogenetic tree of HK proteins clustered all the ComP proteins from various organisms together, instead of gathering them with other HKs of the same organism, indicating an early origin of ComP (Dogsa et al., 2014). The phylogenetic analyses presented here are unrooted and assume an early origin of these pathways in bacterial evolution. We present the distribution and evolutionary relationships of different HK and RR protein types and families, but a larger-scale analysis of all available sequences for two-component QS systems and other HK/RR pairs would be needed to confidently address the issue of the root of quorum sensing.

## Supporting information

Supplementary dataset (accessions, alignments and trees)

## References

Agrawal, R., Sahoo, B.K., Saini, D.K., 2016. Cross-talk and specificity in two-component signal transduction pathways. Future Microbiol. 11, 685–697. https://doi.org/10.2217/fmb-2016-0001

Bandara, H.M.H.N., Lam, O.L.T., Jin, L.J., Samaranayake, L., 2012. Microbial chemical signaling: A current perspective. Crit. Rev. Microbiol. 38, 217–249. https://doi.org/10.3109/1040841X.2011.652065

Brackman, G., Coenye, T., 2014. Quorum Sensing Inhibitors as Anti-Biofilm Agents. Curr. Pharm. Des. 21, 5–11. https://doi.org/10.2174/1381612820666140905114627

Capra, E.J., Laub, M.T., 2012. Evolution of Two-Component Signal Transduction Systems. Annu. Rev. Microbiol. 66, 325–347. https://doi.org/10.1146/annurev-micro-092611-150039

Curtis, M.M., Russell, R., Moreira, C.G., Adebesin, A.M., Wang, C., Williams, N.S., Taussig, R., Stewart, D., Zimmern, P., Lu, B., Prasad, R.N., Zhu, C., Rasko, D.A., Huntley, J.F., Falck, J.R., Sperandio, V., 2014. QseC inhibitors as an antivirulence approach for gram-negative pathogens. MBio 5. https://doi.org/10.1128/mBio.02165-14

Defoirdt, T., 2018. Quorum-Sensing Systems as Targets for Antivirulence Therapy. Trends Microbiol. 26, 313–328. https://doi.org/10.1016/j.tim.2017.10.005

Dereeper, A., Guignon, V., Blanc, G., Audic, S., Buffet, S., Chevenet, F., Dufayard, J.F., Guindon, S., Lefort, V., Lescot, M., Claverie, J.M., Gascuel, O., 2008. Phylogeny.fr: robust phylogenetic analysis for the non-specialist. Nucleic Acids Res. 36, W465–W469. https://doi.org/10.1093/nar/gkn180

Dogsa, I., Choudhary, K.S., Marsetic, Z., Hudaiberdiev, S., Vera, R., Pongor, S., Mandic-Mulec, I., 2014. ComQXPA quorum sensing systems may not be unique to Bacillus subtilis: A census in prokaryotic genomes. PLoS One 9. https://doi.org/10.1371/journal.pone.0096122

Edgar, R.C., 2004. MUSCLE: Multiple sequence alignment with high accuracy and high throughput. Nucleic Acids Res. 32, 1792–1797. https://doi.org/10.1093/nar/gkh340

Forst, S., Kim, D., 2015. Genomic analysis of the histidine kinase family in bacteria and archaea. Microbiology 147, 1197–1212. https://doi.org/10.1099/00221287-147-5-1197

Freedman, J.C., Li, J., Mi, E., McClane, B.A., 2019. Identification of an important orphan histidine kinase for the initiation of sporulation and enterotoxin production by Clostridium perfringens type F strain SM101. MBio 10, 1–17. https://doi.org/10.1128/mBio.02674-18

Galperin, M.Y., 2006. Structural classification of bacterial response regulators: Diversity of output domains and domain combinations. J. Bacteriol. 188, 4169–4182. https://doi.org/10.1128/JB.01887-05

Grebe, T.W., Stock, J.B., 1999. The histidine protein kinase superfamily. Adv. Microb. Physiol. 41, 139–227. https://doi.org/10.1016/s0065-2911(08)60167-8

Hakenbeck, R., Stock, J.B., 1996. Analysis of two-component signal transduction systems involved in transcriptional regulation. Methods Enzymol. 273, 281–300. https://doi.org/10.1016/s0076-6879(96)73026-4

Jacob, S., Foster, A.J., Yemelin, A., Thines, E., 2014. Histidine kinases mediate differentiation, stress response, and pathogenicity in Magnaporthe oryzae. Microbiol. Open 3, 668– 687. https://doi.org/10.1002/mbo3.197

Jin, Y., Li, M., Shang, Y., Liu, L., Shen, X., Lv, Z., Hao, Z., Duan, J., Wu, Y., Chen, C., Pan, J., Yu, F., 2018. Sub-inhibitory concentrations of mupirocin strongly inhibit alpha-toxin production in high-level mupirocin-resistant MRSA by down-regulating agr, saeRS, and sarA. Front. Microbiol. 9, 1–8. https://doi.org/10.3389/fmicb.2018.00993

Jun, S.R., Sims, G.E., Wu, G.A., Kim, S.H., 2010. Whole-proteome phylogeny of prokaryotes by feature frequency profiles: An alignment-free method with optimal feature resolution. Proc. Natl. Acad. Sci. U. S. A. 107, 133–138. https://doi.org/10.1073/pnas.0913033107

Kaserer, A.O., West, A.H., 2010. Histidine kinases in two-component signaling pathways, in: Handbook of Cell Signaling, 2/E. Elsevier Inc., pp. 581–586. https://doi.org/10.1016/B978-0-12-374145-5.00078-4

Kaur, A., Capalash, N., Sharma, P., 2018. Quorum sensing in thermophilesl: prevalence of autoinducer-2 system. BMC Microbiol. 18, 62.

Koretke, K.K., Lupas, A.N., Warren, P. V, Rosenberg, M., Brown, J.R., 2000. Evolution of Two-Component Signal Transduction. Mol Biol Evol. 17, 1956–1970.

Koretke, K.K., Volker, C., Bower, M.L., Lupas, A.N., 2003. Molecular Evolution of Histidine Kinases. Histidine Kinases Signal Transduct. 483–506. https://doi.org/10.1016/B978-012372484-7/50024-2

Lerat, E., Moran, N.A., 2004. The Evolutionary History of Quorum-Sensing Systems in Bacteria. Mol. Biol. Evol. 21, 903–913. https://doi.org/10.1093/molbev/msh097

Liu, C., Worthington, R.J., Melander, C., Wu, H., 2011. A new small molecule specifically inhibits the cariogenic bacterium Streptococcus mutans in multispecies biofilms. Antimicrob. Agents Chemother. 55, 2679–2687. https://doi.org/10.1128/AAC.01496-10

Madeira, F., Park, Y.M., Lee, J., Buso, N., Gur, T., Madhusoodanan, N., Basutkar, P., Tivey, A.R.N., Potter, S.C., Finn, R.D., Lopez, R., 2019. The EMBL-EBI search and sequence analysis tools APIs in 2019. Nucleic Acids Res. 47, W636–W641. https://doi.org/10.1093/nar/gkz268

Mascher, T., Helmann, J.D., Unden, G., 2006. Stimulus Perception in Bacterial Signal-Transducing Histidine Kinases. Microbiol. Mol. Biol. Rev. 70, 910–938. https://doi.org/10.1128/mmbr.00020-06

Ng, W.-L., Bassler, B.L., 2009. Bacterial Quorum-Sensing Network Architectures. Annu. Rev. Genet. 43, 197–222. https://doi.org/10.1146/annurev-genet-102108-134304

Papenfort, K., Bassler, B.L., 2016. Quorum sensing signal-response systems in Gram-negative bacteria. Nat. Rev. Microbiol. 14, 576–588. https://doi.org/10.1038/nrmicro.2016.89

Pearson, W.R., 2013. Selecting the Right Similarity-Scoring Matrix. Curr Protoc Bioinforma. 43, 3.5.1–3.5.9. https://doi.org/10.1002/0471250953.bi0305s43

Rasmussen, B.B., Nielsen, K.F., Machado, H., Melchiorsen, J., Gram, L., Sonnenschein, E.C., 2014. Global and Phylogenetic Distribution of Quorum Sensing Signals, Acyl Homoserine Lactones, in the Family of Vibrionaceae. Mar. Drugs 12, 5527–5546. https://doi.org/10.3390/md12115527

Reading, N.C., Sperandio, V., 2006. Quorum sensing: The many languages of bacteria. FEMS Microbiol. Lett. 254, 1–11. https://doi.org/10.1111/j.1574-6968.2005.00001.x

Rutherford, S.T., Bassler, B.L., 2012. Bacterial quorum sensing: Its role in virulence and possibilities for its control. Cold Spring Harb. Perspect. Med. 2:a012427. https://doi.org/10.1101/cshperspect.a012427

Scoffone, V.C., Trespidi, G., Chiarelli, L.R., Barbieri, G., Buroni, S., 2019. Quorum sensing as antivirulence target in cystic fibrosis pathogens. Int. J. Mol. Sci. 20. https://doi.org/10.3390/ijms20081838

Sreenivasulu, Y., 2015. THE POTENTIAL OF MICROBIAL WEALTH FOR PHYTOPATHOGEN MANAGEMENT AND PLANT GROWTH ENHANCEMENT. J. Biol. Nat. 2(1), 6–15.

Stamatakis, A., 2014. RAxML version 8: A tool for phylogenetic analysis and post-analysis of large phylogenies. Bioinformatics 30, 1312–1313. https://doi.org/10.1093/bioinformatics/btu033

Sully, E.K., Malachowa, N., Elmore, B.O., Alexander, S.M., Femling, J.K., Gray, B.M., DeLeo, F.R., Otto, M., Cheung, A.L., Edwards, B.S., Sklar, L.A., Horswill, A.R., Hall, P.R., Gresham, H.D., 2014. Selective Chemical Inhibition of agr Quorum Sensing in Staphylococcus aureus Promotes Host Defense with Minimal Impact on Resistance. PLoS Pathog. 10. https://doi.org/10.1371/journal.ppat.1004174

Tiwari, R., Karthik, K., Rana, R., Malik, Y.S., Dhama, K., Joshi, S.K., 2016. Quorum sensing inhibitors/antagonists countering food spoilage bacteria-need molecular and pharmaceutical intervention for protecting current issues of food safety. Int. J. Pharmacol. 12, 262–271. https://doi.org/10.3923/ijp.2016.262.271

Wang, W., Shu, D., Chen, L., Jiang, W., Lu, Y., 2009. Cross-talk between an orphan response regulator and a noncognate histidine kinase in streptomyces coelicolor. FEMS Microbiol. Lett. 294, 150–156. https://doi.org/10.1111/j.1574-6968.2009.01563.x

West, A.H., Stock, A.M., 2001. Histidine kinases and response regulator proteins in two-component signaling systems. Trends Biochem. Sci. 26, 369–376. https://doi.org/10.1016/S0968-0004(01)01852-7

Whiteley, M., Diggle, S.P., Greenberg, E.P., 2017. Progress in and promise of bacterial quorum sensing research. Nature 551, 313–320. https://doi.org/10.1038/nature24624

Wilke, K.E., Francis, S., Carlson, E.E., 2015. Inactivation of multiple bacterial histidine kinases by targeting the ATP-binding domain. ACS Chem. Biol. 10, 328–335. https://doi.org/10.1021/cb5008019

Worthington, R.J., Blackledge, M.S., Melander, C., 2013. Small-molecule inhibition of bacterial two-component systems to combat antibiotic resistance and virulence. Future Med. Chem. 5, 1265–1284. https://doi.org/10.4155/fmc.13.58

