## Supplementary Figures and Tables for "Evolution of two-component quorum sensing systems"

Supplementary Figures and Supplementary Tables

Supplementary Figure S1: Classification of the HKs-RRs cognate pairs in the KEGG database (KEGG map: ko2020)

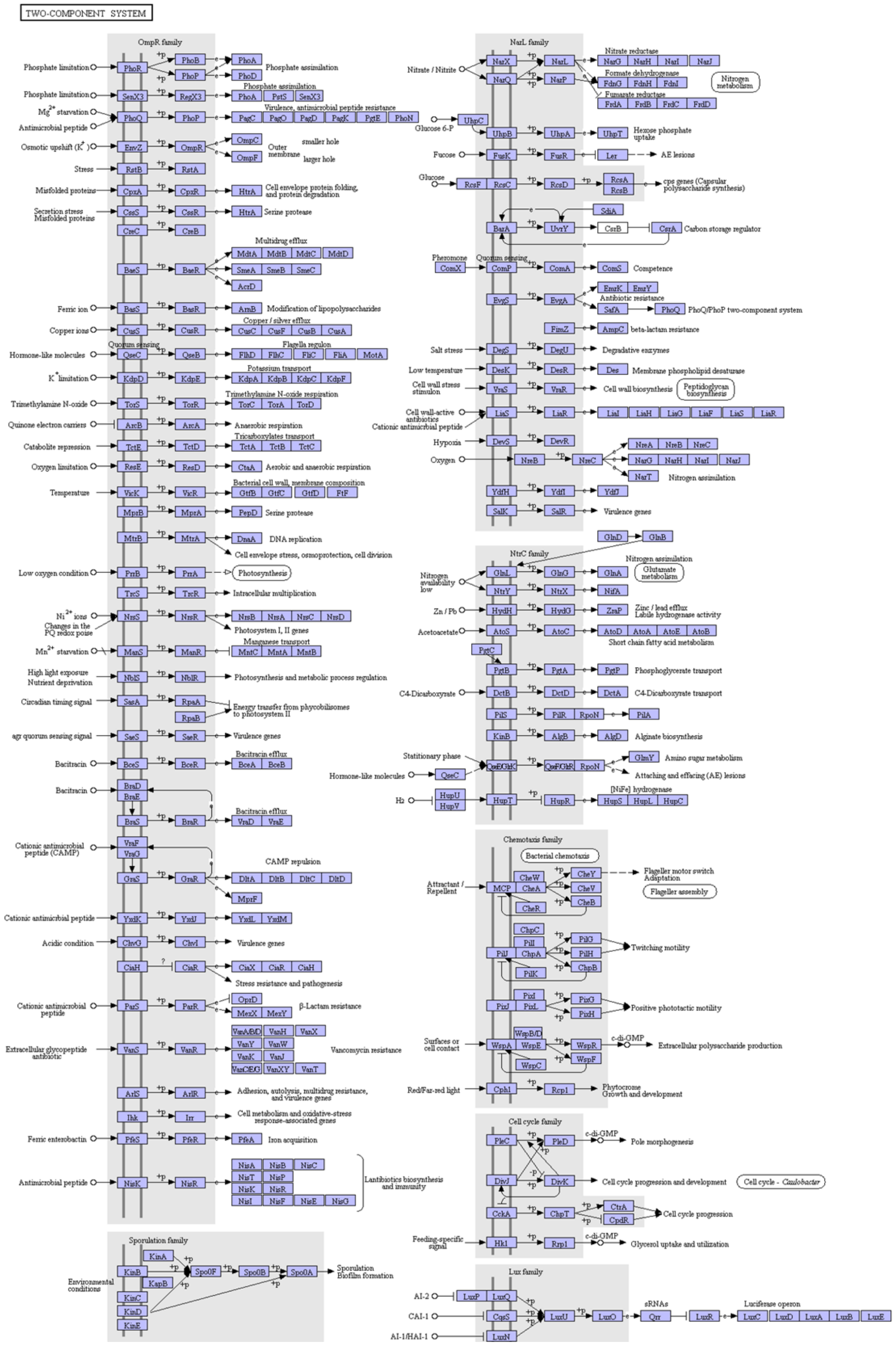

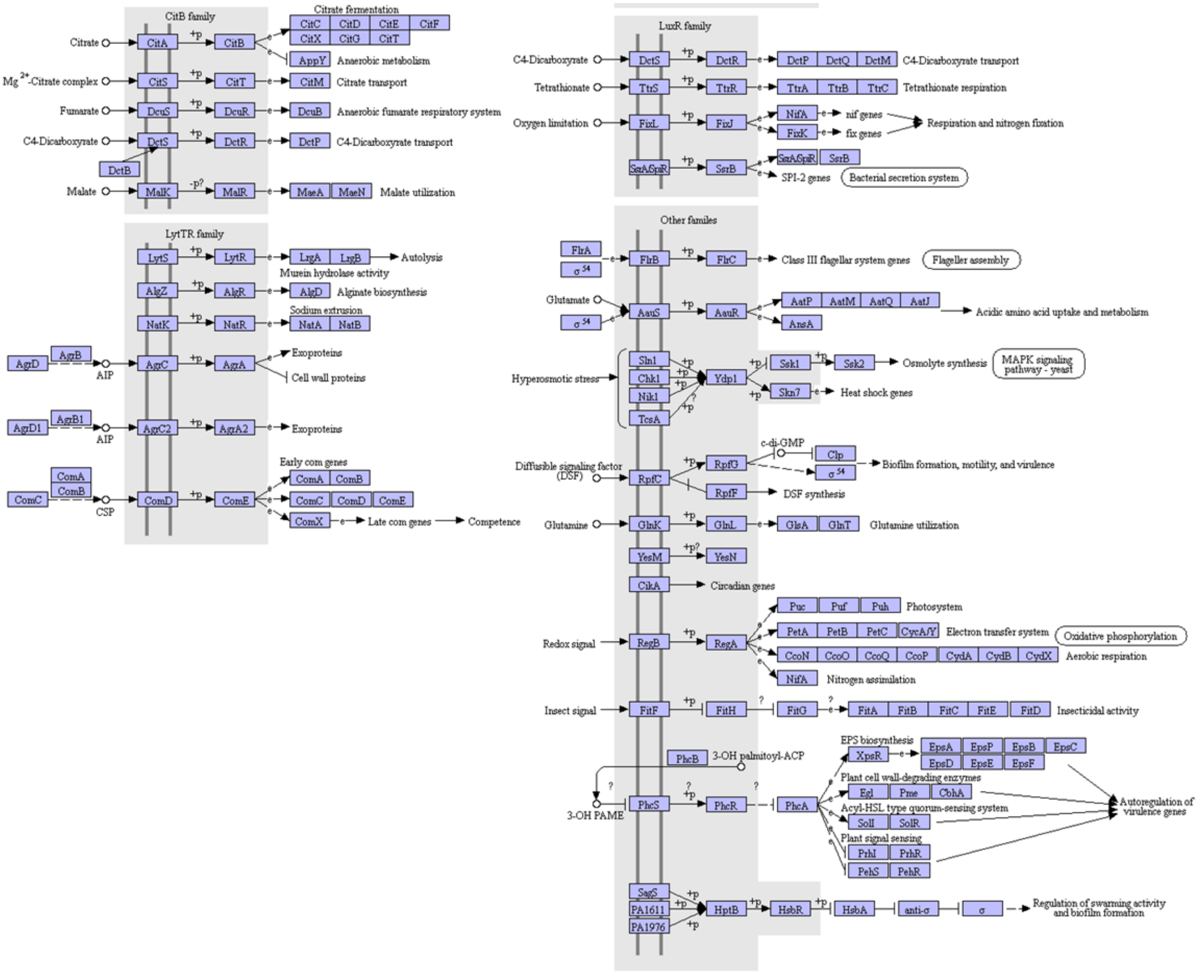

02020 7/9/19
(c) Kanehisa Laboratories

Supplementary Figure S2A: MAFFT alignment of all the HKs

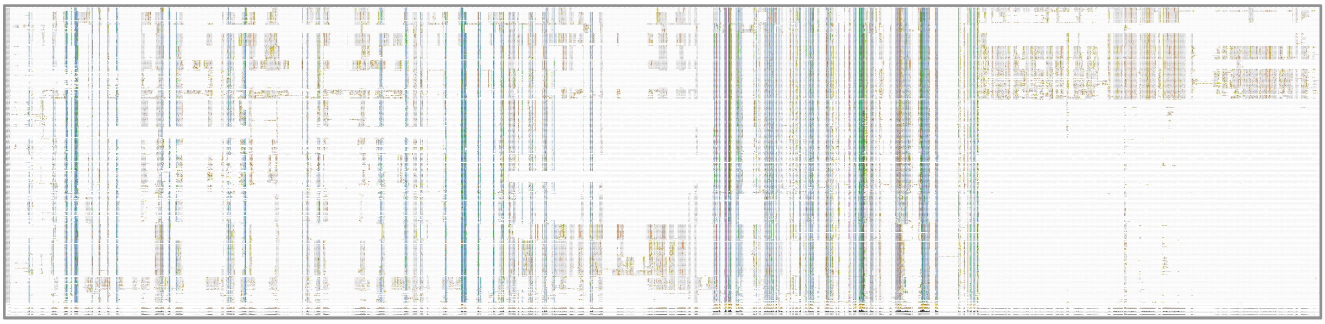

Supplementary Figure S2B: The parts of the alignment in grey were removed using Mesquite.

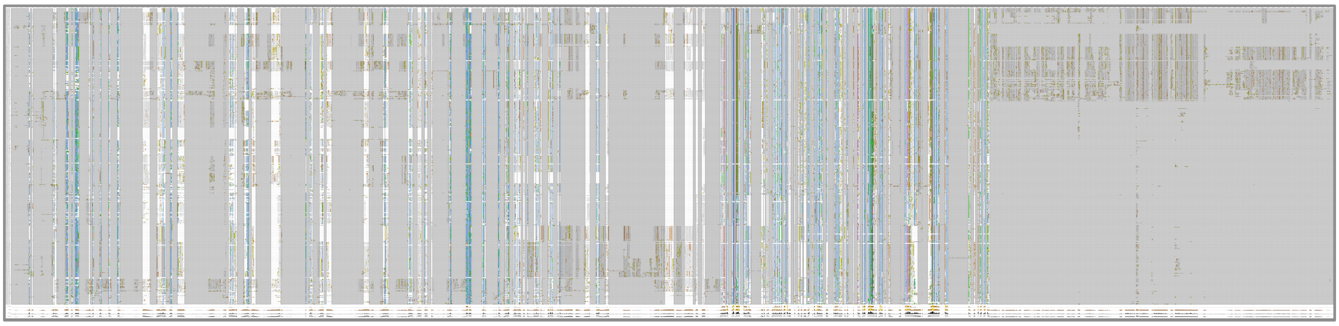

Supplementary Figure S2C: The final part of the MAFFT HKs alignment that remained after the trimming.

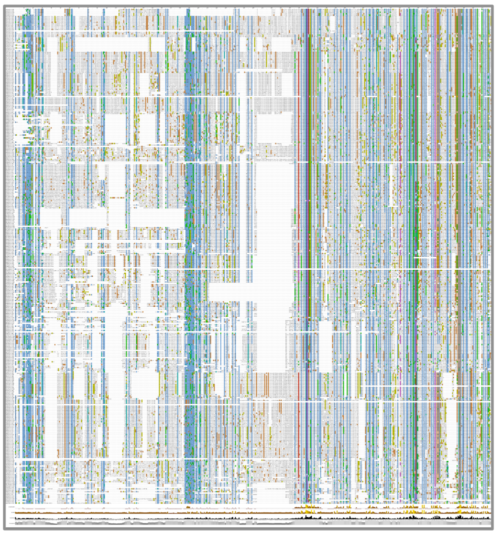

Supplementary Figure S3A: MUSCLE alignment of all the HKs

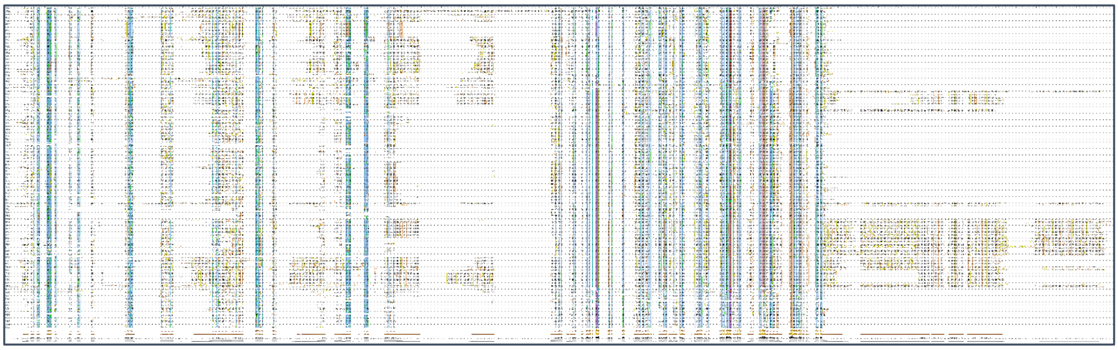

Supplementary Figure S3B: The parts of the alignment in grey were removed using Mesquite.

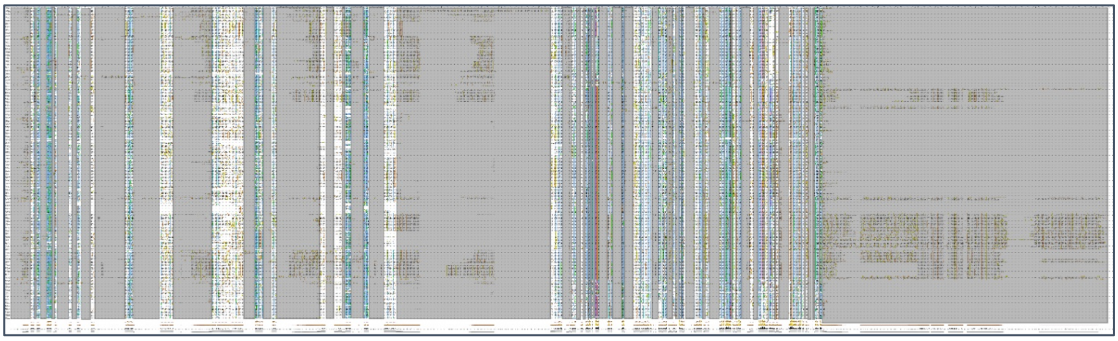

Supplementary Figure S3C: The final part of the MUSCLE HKs alignment that remained after the trimming.

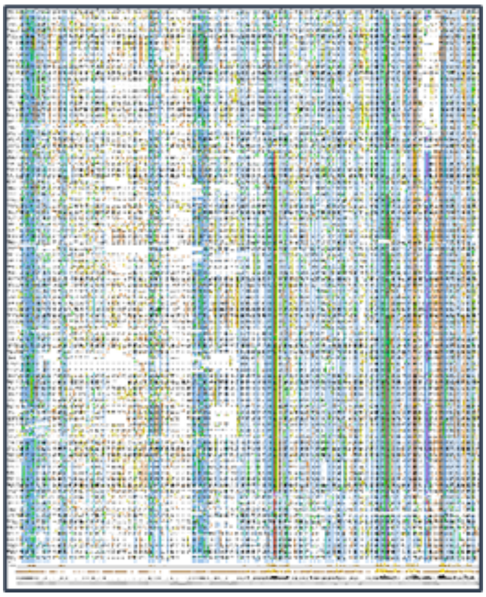

Supplementary Figure S4A: The MAFFT alignment of all the RRs.

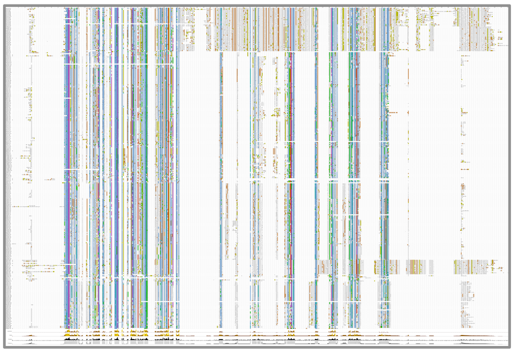

Supplementary Figure S4B: The parts of the alignment in grey were removed using Mesquite.

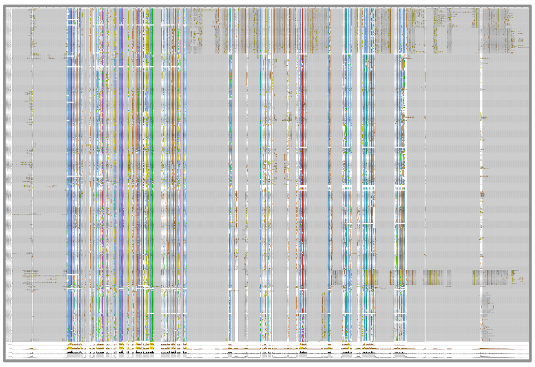

Supplementary Figure S4C: The final part of the MAFFT RRs alignment that remained after the trimming.

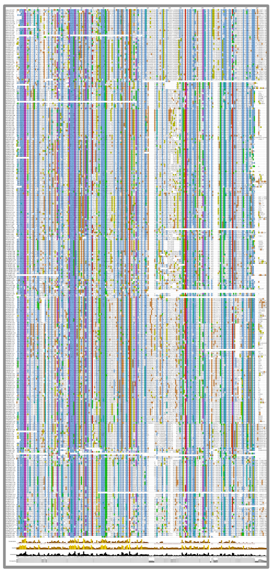

Supplementary Figure S5A: MUSCLE alignment of all the RRs.

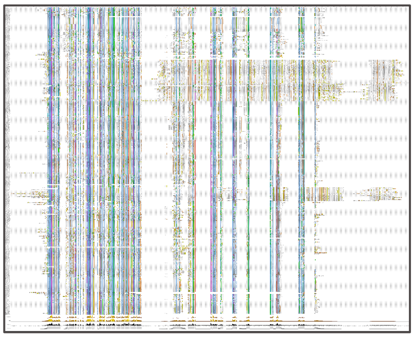

Supplementary Figure S5B: The parts of the alignment in grey were removed using Mesquite.

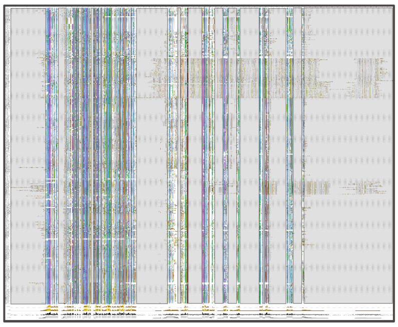

Supplementary Figure S5C: The final part of the MUSCLE RRs alignment that remained after the trimming.

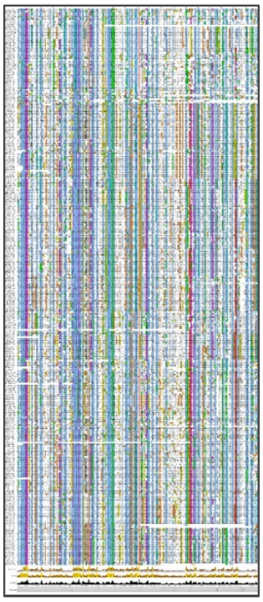

Supplementary Figure S6: Phylogenetic analysis of the NarL family. Both trees reconstructed using PhyML based on the MUSCLE alignment. (A) Histidine Kinases. (B) Response Regulators. The color of the bullets indicates the bootstrap value of the nodes. White: 50% -80%, Grey: 80%-95%, Black: over 95%

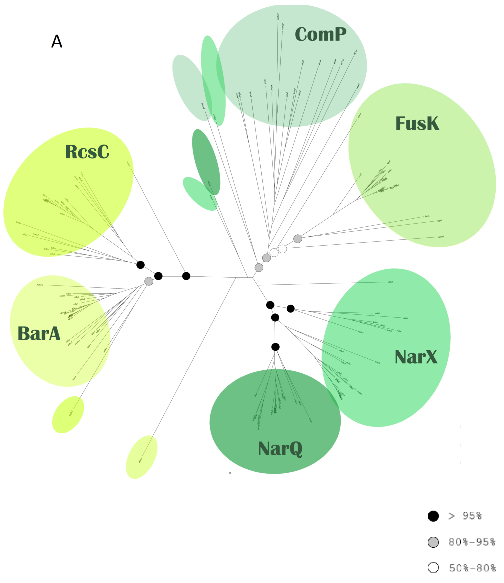

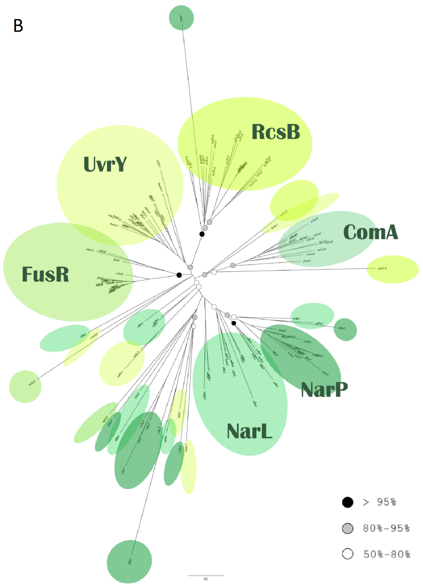

Supplementary Figure S7: Phylogenetic analysis of the OmpR family. Both trees reconstructed using PhyML based on the MUSCLE alignment. (A) Histidine Kinases. (B) Response Regulators.

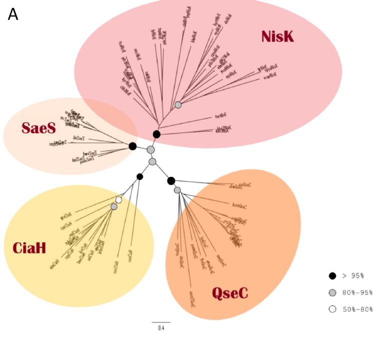

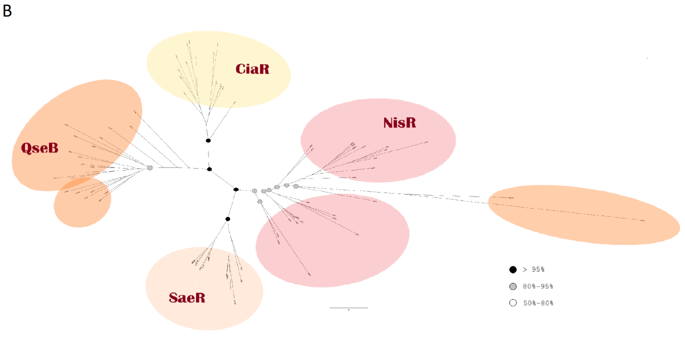

Supplementary Figure S8: Phylogenetic analysis of the Lux family histidine kinases. Tree reconstructed using PhyML based on the MUSCLE alignment.

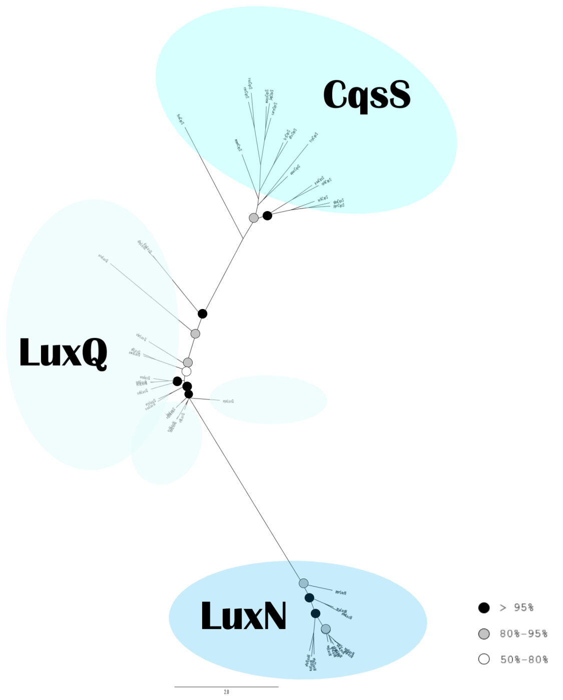

Supplementary Figure S9: Phylogenetic analysis of the LytR family. (A) Histidine Kinases. (B) Response Regulators. Both trees reconstructed using PhyML based on the MUSCLE alignment.

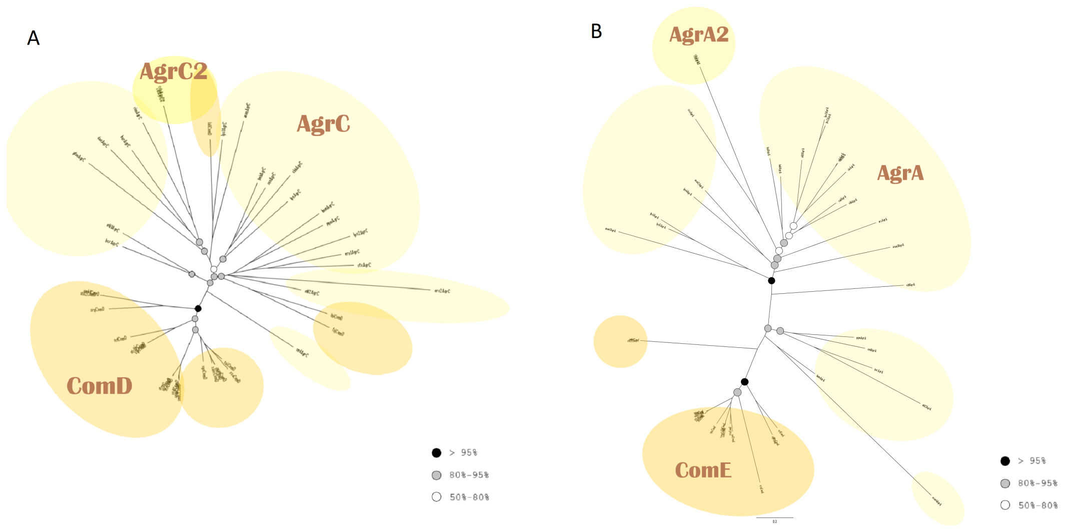

Supplementary Figure S10: Phylogenetic analysis of all histidine kinase families. Tree reconstructed using RaxML based on the MUSCLE alignment.

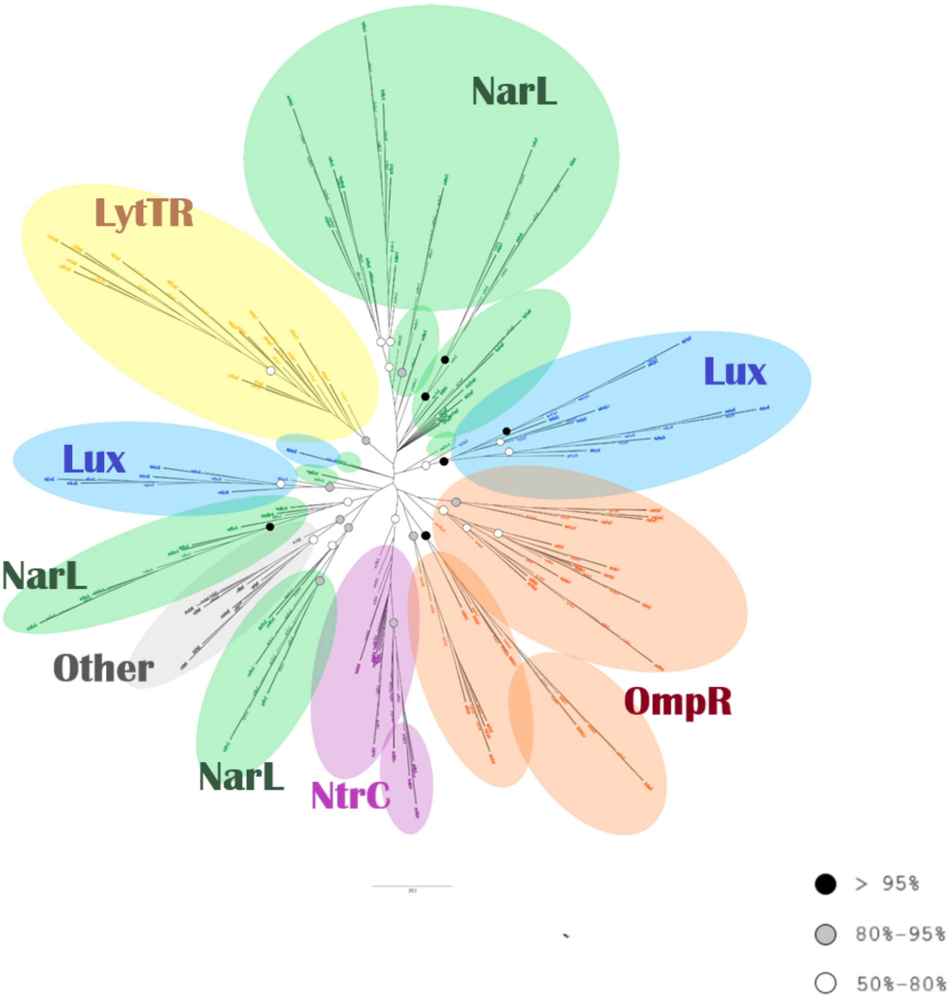

Supplementary Figure S11: Phylogenetic analysis of all response regulator families. Tree reconstructed using RaxML based on the MUSCLE alignment

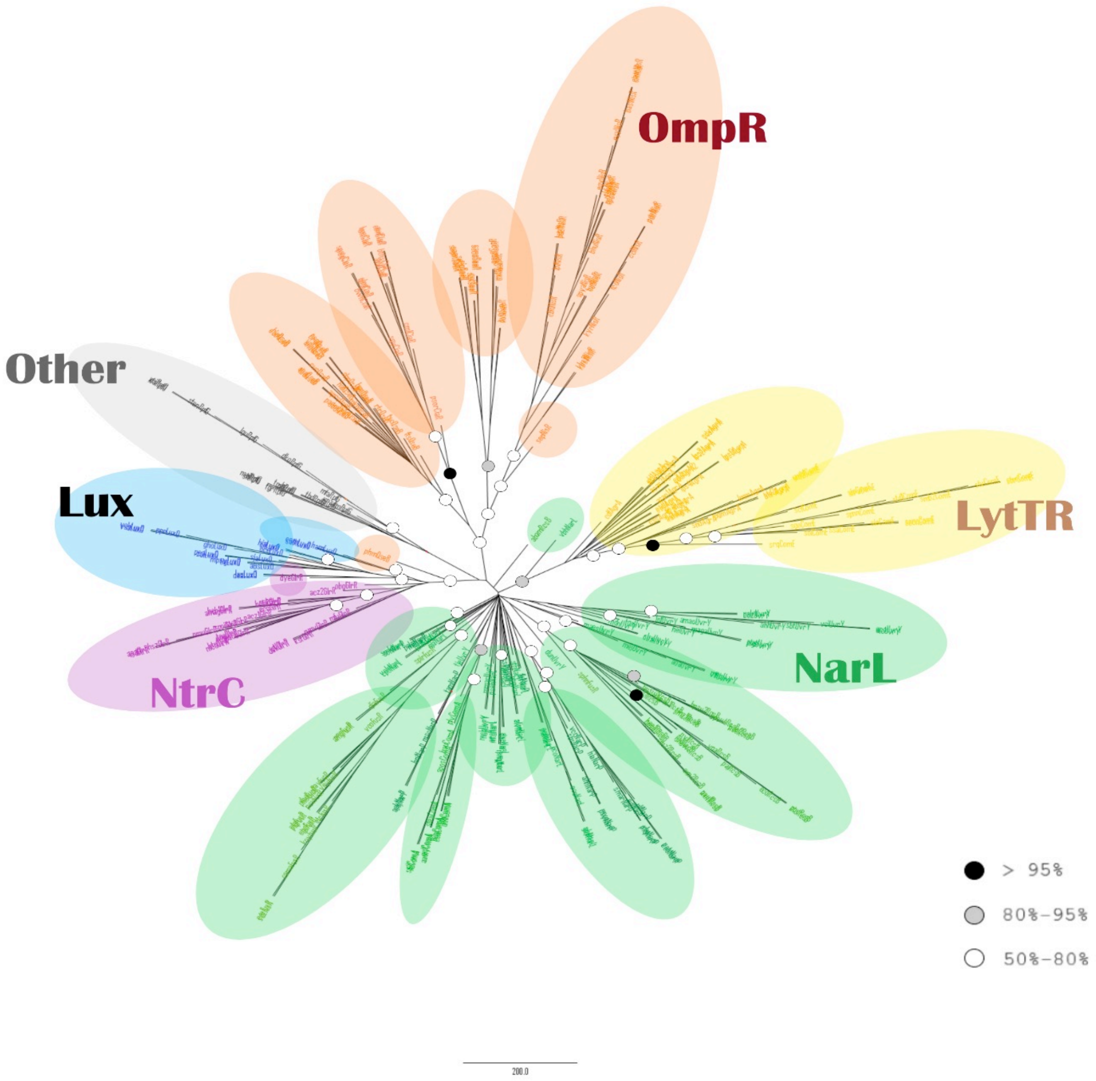

Supplementary Table S1: PFAM and InterPro entries for each domain discussed in this study, as well as the entry of the superfamily (NCBI-CDD) and Pfam clan they belong to.

| ***DOMAINS*** | **PFAM entry** | **InterPro entry** | **superfamily(NCBI-CDD)** | **pfam clan** |
|---|---|---|---|---|
| *LuxQ-periplasm* | PF09308 | IPR015387 | cl07810 | CL0165 |
| *Response_reg* | PF00072 | IPR001789 | cl19078 | CL0304 |
| *PilJ* | PF13675 | IPR029095 | cl16344 | … |
| *HAMP* | PF00672 | IPR003660 | cl01054 | CL0681 |
| *HisKA* | PF00512 | IPR003661 | cl29579 | CL0025 |
| *HisKA_3* | PF07730 | IPR011712 | cl38245 | |
| *HATPase_c* | PF02518 | IPR003594 | cl00075 | |
| *HATPase_c_2* | PF13581 | IPR003594 | | |
| *HATPase_c_5* | PF14501 | IPR032834 | | |
| *RcsC* | PF09456 | IPR019017 | cl09693 | CL0304 |
| *DUF2222* | PF09984 | IPR019247 | cl10814 | CL0165 |
| *Hpt* | PF01627 | IPR008207 | cl00086 | … |
| *2CSK_N* | PF08521 | IPR013727 | cl07224 | CL0165 |
| *Peripla_BP_4* | PF13407 | IPR025997 | cl10011 | CL0144 |
| *Sigma54_activat* | PF0158 | IPR002078 | cl37971 | CL0023 |
| *Sigma54_activ_2* | PF14532 | IPR002078 | cl37881 | |
| *Sigma54_AID* | PF00309 | IPR000394 | cl08259 | … |
| *Sigma54_DBD* | PF04552 | IPR007634 | cl10632 | CL0123 |
| *Sigma54_CBD* | PF04963 | IPR007046 | cl29309 | |
| *Sigma70_r4_2* | PF08281 | IPR013249 | cl22867 | |
| *Sigma70_r4* | PF04545 | IPR007630 | | |
| *Trans_reg_C* | PF00486 | IPR001867 | cl21459 | |
| *AAA_5* | PF07728 | IPR011704 | cl38244 | CL0023 |
| *HTH_8* | PF02954 | IPR002197 | cl22895 | |
| *GerE* | PF00196 | IPR000792 | cl21459 | |
| *RcsD_ABL* | PF16359 | IPR032306 | cl24808 | CL0304 |
| *HD* | PF01966 | IPR006674 | cl21469 | CL0237 |
| *HD_5* | PF13487 | … | | |
| *ECH_1* | PF00378 | IPR001753 | cl23717 | CL0127 |
| *ECH_2* | PF16113 | IPR032259 | | |
| *MASE1* | PF05231 | IPR007895 | … | … |
| *LytTR* | PF04397 | IPR007492 | cl04498 | CL0049 |
| *RcsF* | PF16358 | IPR030852 | cl08147 | CL0522 |

Supplementary Table S2: The presence of the proteins in several QS pathways was examined in various bacterial classsifications.

| Phylum | Class | Family | Genus |
|---|---|---|---|
| gemmatimonadetes | a-proteobacteria | aquificae | leptospira (spirocahetes) |
| acidobacteria | b-proteobacteria | | deinococci |
| chlorobi | g-proteobacteria | | nitrospira |
| bacteroidetes | d-proteobacteria | | chlamydiae |
| verrucomicrobia | e-proteobacteria | | borrelia (spirochaetes) |
| chloroflexi | clostridia (firmicutes) | | |
| cyanobacteria | bacilli (firmicutes) | | |
| actinobacteria | | | |
| fusobacteria | | | |
| mollicutes (tenericutes) | | | |
| dictyoglomi | | | |
| thermotogae | | | |
| elusimicrobia | | | |
| Planctomycetes | | | |
| Fibrobacteres | | | |
| Chrysiogenetes | | | |
| Deferribacteres | | | |
| Other Spirochaetes | | | |

Supplementary Table S3A: The number of HK amino acid sequences which were used for the phylogenetic trees.

| | NarL family | | | OmpR family | | | Lux family | | | LytTR family | | NtrC family | | |
|---|---|---|---|---|---|---|---|---|---|---|---|---|---|---|
| | Taxa | Families | Total | Taxa | Families | Total | Taxa | Families | Total | Families | Total | Taxa | Families | Total |
| | NarX | | | | QseC | | LuxQ | | | AgrC | | GlrK (QseE) | | |
| *KEGG* | 23 | 50 | 757 | 28 | 68 | 1190 | 3 | 3 | 59 | 24 | 714 | 19 | 44 | 652 |
| *Our study* | 9 | 15 | 25 | 9 | 14 | 20 | 3 | 3 | 17 | 15 | 20 | 16 | 29 | 33 |
| *%* | 39% | 30% | 3% | 32% | 21% | 2% | 100% | 100% | 29% | 63% | 3% | 84% | 66% | 5% |
| | NarQ | | | | SaeS | | LuxN | | | AgrC2 | | | | |
| *KEGG* | 7 | 15 | 476 | 3 | 6 | 169 | 1 | 1 | 35 | 1 | 2 | | | |
| *Our study* | 5 | 10 | 20 | 2 | 6 | 13 | 1 | 1 | 13 | 1 | 2 | | Other | |
| *%* | 71% | 67% | 4% | 67% | 100% | 8% | 100% | 100% | 37% | 100% | 100% | Taxa | Families | Total |
| | BarA | | | | CiaH | | CqsS | | | ComD | | RpfC | | |
| *KEGG* | 20 | 42 | 863 | 5 | 13 | 256 | 7 | 9 | 100% | 2 | 52 | 13 | 19 | 99 |
| *Our study* | 11 | 20 | 23 | 5 | 13 | 14 | 7 | 9 | 100% | 2 | 22 | 13 | 19 | 20 |
| *%* | 55% | 48% | 3% | 100% | 100% | 5% | 100% | 100% | 21% | 100% | 42% | 100% | 100% | 20% |
| | ComP | | | | NisK | | | | | | | | | |
| *KEGG* | 5 | 9 | 159 | 11 | 23 | 367 | | | | | | | | |
| *Our study* | 5 | 9 | 17 | 11 | 21 | 29 | | | | | | | | |
| *%* | 100% | 100% | 11% | 100% | 91% | 8% | | | | | | | | |
| | FusK | | | | | | | | | | | | | |
| *KEGG* | 7 | 12 | 144 | | | | | | | | | | | |
| *Our study* | 12 | 12 | 20 | | | | | | | | | | | |
| *%* | 86% | 100% | 14% | | | | | | | | | | | |
| | RcsC | | | | | | | | | | | | | |
| *KEGG* | 8 | 17 | 781 | | | | | | | | | | | |
| *Our study* | 8 | 17 | 22 | | | | | | | | | | | |
| *%* | 100% | 100% | 3% | | | | | | | | | | | |

Supplementary Table S3B: The number of RR amino acid sequences which were used for the phylogenetic trees.

| | NarL family | | | OmpR family | | | Lux family | | | LytTR family | | | NtrC family | | |
|---|---|---|---|---|---|---|---|---|---|---|---|---|---|---|---|
| | *Taxa* | *Families* | *Total* | *Taxa* | *Families* | *Total* | *Taxa* | *Families* | *Total* | *Taxa* | *Families* | *Total* | *Taxa* | *Families* | *Total* |
| | NarL | | | QseB | | | LuxO | | | AgrA | | | GlrR (QseF) | | |
| *KEGG* | 55 | 85 | 1051 | 36 | 73 | 1386 | 11 | 14 | 129 | 6 | 19 | 623 | 25 | 53 | 683 |
| *Our study* | 17 | 26 | 26 | 23 | 23 | 23 | 11 | 14 | 18 | 6 | 19 | 21 | 24 | 21 | 24 |
| *%* | 31% | 31% | 2% | 64% | 32% | 2% | 100% | 100% | 14% | 100% | 100% | 3% | 96% | 40% | 4% |
| | NarP | | | SaeR | | | | | | AgrA2 | | | | | |
| *KEGG* | 12 | 24 | 503 | 2 | 5 | 156 | | | | 1 | 2 | 2 | | | |
| *Our study* | 12 | 23 | 23 | 2 | 5 | 13 | | | | 1 | 2 | 2 | Other | | |
| *%* | 100% | 96% | 5% | 100% | 100% | 8% | | | | 100% | 100% | 100% | *Taxa* | *Families* | *Total* |
| | UvrY | | | CiaR | | | | | | ComE | | | RpfG | | |
| *KEGG* | 26 | 62 | 924 | 4 | 14 | 261 | | | | 1 | 1 | 49 | 7 | 11 | 83 |
| *Our study* | 22 | 27 | 27 | 4 | 14 | 15 | | | | 1 | 1 | 1 | 7 | 11 | 15 |
| *%* | 85% | 44% | 3% | 100% | 100% | 6% | | | | 100% | 100% | 2% | 100% | 100% | 18% |
| | ComA | | | NisR | | | | | | | | | | | |
| *KEGG* | 2 | 4 | 150 | 10 | 20 | 377 | | | | | | | | | |
| *Our study* | 2 | 4 | 10 | 10 | 20 | 25 | | | | | | | | | |
| *%* | 100% | 100% | 7% | 100% | 100% | 7% | | | | | | | | | |
| | FusR | | | | | | | | | | | | | | |
| *KEGG* | 6 | 12 | 193 | | | | | | | | | | | | |
| *Our study* | 6 | 12 | 20 | | | | | | | | | | | | |
| *%* | 100% | 100% | 10% | | | | | | | | | | | | |
| | RcsB | | | | | | | | | | | | | | |
| *KEGG* | 8 | 17 | 1241 | | | | | | | | | | | | |
| *Our study* | 8 | 14 | 24 | | | | | | | | | | | | |
| *%* | 100% | 82% | 2% | | | | | | | | | | | | |
